# Species boundaries and molecular markers for the classification of 16SrI phytoplasmas inferred by genome analysis

**DOI:** 10.1101/2020.01.31.928135

**Authors:** Shu-Ting Cho, Hung-Jui Kung, Weijie Huang, Saskia A. Hogenhout, Chih-Horng Kuo

## Abstract

Phytoplasmas are plant-pathogenic bacteria that impact agriculture worldwide. The commonly adopted classification system for phytoplasmas is based on the restriction fragment length polymorphism (RFLP) analysis of their 16S rRNA genes. With the increased availability of phytoplasma genome sequences, the classification system can now be refined. This work examined 11 strains in the 16SrI group within the genus ‘*Candidatus* Phytoplasma’ and investigated the possible species boundaries. We confirmed that the RFLP classification method is problematic due to intragenomic variation of the 16S rRNA genes and uneven weighing of different nucleotide positions. Importantly, our results based on the molecular phylogeny, differentiations in chromosomal segments and gene content, and divergence in homologous sequences, all supported that these strains may be classified into multiple operational taxonomic units (OTUs) equivalent to species. Strains assigned to the same OTU share >97% genome-wide average nucleotide identity (ANI) and >78% of their protein-coding genes. In comparison, strains assigned to different OTUs share <94% ANI and <75% of their genes. Reduction in homologous recombination between OTUs is one possible explanation for the discontinuity in genome similarities, and these findings supported the proposal that 95% ANI could serve as a cutoff for distinguishing species in bacteria. Additionally, critical examination of these results and the raw sequencing reads led to the identification of one genome that was presumably mis-assembled by combining two sequencing libraries built from phytoplasmas belonging to different OTUs. This finding provided a cautionary tale for working on uncultivated bacteria. Based on the new understanding of phytoplasma divergence and the current genome availability, we developed five molecular markers that could be used for multilocus sequence analysis (MLSA). By selecting markers that are short yet highly informative, and are distributed evenly across the chromosome, these markers provided a cost-effective system that is robust against recombination. Finally, examination of the effector gene distribution further confirmed the rapid gains and losses of these genes, as well as the involvement of potential mobile units (PMUs) in their molecular evolution. Future improvements on the taxon sampling of phytoplasma genomes will allow further expansions of similar analysis, and thus contribute to phytoplasma taxonomy and diagnostics.

## Introduction

Phytoplasmas are a group of insect-transmitted plant-pathogenic bacteria that reduce yields of diverse crops worldwide (Lee et al., 2000; Hogenhout et al., 2008; Bertaccini and Lee, 2018). These obligate parasites are related to the animal-pathogenic mycoplasmas (Chen et al., 2012), and both groups of these wall-less bacteria were assigned to the class Mollicutes under the phylum Tenericutes (Brown, 2010). However, unlike mycoplasmas and several other Mollicutes lineages, pure culture of phytoplasmas outside of their hosts has not been achieved yet. Due to this reason, all phytoplasmas were assigned to a ‘*Candidatus*’ (‘*Ca*.’) genus (The IRPCM Phytoplasma/Spiroplasma Working Team, 2004).

For the purposes of pathogen detection or research on their basic biology, a reliable system for strain identification and classification is critical. For example, it is important to understand if the diseases affecting different plants, transmitted by different vectors, and/or occurring in different locations are caused by the same or different phytoplasma strains. In addition, to infer the evolutionary history and biodiversity of phytoplasmas, defining species or other equivalent biological units are of fundamental importance. The commonly used system for phytoplasma classification was first established in the 1990s (Lee et al., 1993; Lee et al., 1998). This system utilizes a defined set of 17 restriction enzymes to perform restriction fragment length polymorphism (RFLP) analysis on a 1.25-kb PCR product derived from their 16S rRNA gene (Gundersen and Lee, 1996). Under this system, a set of reference strains were chosen, and a new strain could be assigned to the same 16S rRNA gene RFLP (16Sr) group and subgroup as a reference when the similarity coefficient is higher than 0.85 and 0.97, respectively (Zhao et al., 2009; Zhao and Davis, 2016). With the development of an easy-to-use web-based tool *i*PhyClassifier (Zhao et al., 2009), this system has been well-adopted by the phytoplasma research community.

However, despite the popularity, this 16Sr classification system has several shortcomings. To begin with, the RFLP approach considers only those nucleotides located within the restriction sites, rather than the entire sequence. Additionally, the similarity coefficients of the RFLP patterns do not provide information about phylogenetic relationships. Furthermore, intragenomic sequence variations of 16S rRNA genes may cause problems in classification. Most importantly, although 16S rRNA gene is considered as a universal marker for bacterial classification, the use of a single locus is inherently unreliable. Rather, adoption of multilocus sequence analysis (MLSA) is strongly recommended (Glaeser and Kämpfer, 2015). Unfortunately, selection of suitable markers for MLSA is not straightforward. Due to the highly variable evolutionary rates among different bacteria and among different genes (Kuo and Ochman, 2009), the markers that work well for one group do not necessary work well for others. Although extensive efforts have been devoted to the development of MLSA markers for phytoplasmas (Martini et al., 2019), genome-level assessment for the relative performance of these markers is lacking. Moreover, while the use of genome analysis to replace MLSA provides a potential solution and is gaining popularity (Konstantinidis and Tiedje, 2005; Konstantinidis et al., 2017; Jain et al., 2018; Parks et al., 2018), genome analysis is still challenging for uncultured bacteria such as phytoplasmas. For these reasons, development and evaluation of MLSA markers are still important.

To develop and evaluate MLSA markers, the availability of genome sequences provides a comprehensive guide for PCR primer design. For example, based on the complete genome sequence of ‘*Candidatus* Phytoplasma asteris’ OY-M (Oshima et al, 2004), 18 PCR primer sets were designed to cover different regions of the chromosome (Kakizawa and Kamagata, 2014). Moreover, by developing a multiplex-PCR method for these primer sets and establishing the amplification patterns in a collection of reference strains, this previous study provided a fast and cost-effective method for strain identification (Kakizawa and Kamagata, 2014). As the number of available genome sequences increases, the genome-enabled MLSA maker development could be further improved in two critical aspects. First, the genome-scale molecular phylogeny provides a reliable reference for evaluating the performance of each candidate markers, such that the best-performing ones could be selected. Second, with the comprehensive genomic information of all reference strains, the chromosomal location of each candidate markers and the exact primer sequences could be evaluated using bioinformatic methods. In this regard, phytoplasmas belonging to the 16SrI group, also often known as the aster yellows (AY) group or ‘*Ca.* P. asteris’ (i.e., a provisional species defined as encompassing all known subgroups within the 16SrI group) (Lee et al., 2004), represent the best system to test the concept of genome-assisted MLSA marker development. Among the >30 16Sr groups that have been described (Zhao and Davis, 2016), only eight have genome sequences available (Cho et al., 2019b). Moreover, the genome sequencing efforts have been highly focused on the 16SrI group (Table 1), which has a worldwide geographical distribution, wide range of plant hosts, and great impact on agriculture (Lee et al., 2004). Finally, this group is also the one that received most research attention on the molecular mechanisms of plant-microbe interactions; most of the phytoplasma effectors that have been characterized were initially studied in 16SrI phytoplasmas (Bai et al., 2009; Hoshi et al., 2009; MacLean et al., 2011; Wang et al. 2018b; Huang and Hogenhout, 2019). Previous studies of this group identified extensive levels of genome divergence and suggested that ‘*Ca.* P. asteris’ may be classified into multiple operational taxonomic units (OTUs) equivalent to species (Firrao et al., 2013; Cho et al., 2019a). In this work, we expanded the sampling of available genome sequences to perform in-depth analysis of genome comparisons. Specifically, we aimed to provide quantitative guidelines for defining putative species boundaries, which could better inform phytoplasma taxonomy. Additionally, we aimed to develop MLSA markers that could facilitate future genetic characterization of the phytoplasmas belonging to the 16SrI or AY group.

**Table 1.** List of the genome sequences analyzed.

| Strain | Accession | Sequencing platform and depth | Assembly | Genome Size (bp) | CDS-intact | CDS-pseudo | Location | Host | Reference |
| --- | --- | --- | --- | --- | --- | --- | --- | --- | --- |
| AYWB | GCF_000012225 | Sanger; not available | Complete | 723,970 | 557 | 116 | USA | Lettuce | Bai et al., 2006 |
| NJAY | GCA_002554195 | Roche/454; 200x | 75 contigs | 652,092 | 371 | 262 | USA | Lettuce | Sparks et al., 2018 |
| WBD | GCF_000495255 | Illumina; 5,000x | 6 contigs | 611,462 | 471 | 76 | China | Wheat | Chen et al., 2014 |
| OY-M | GCF_000009845 | Sanger; not available | Complete | 853,092 | 728 | 144 | Japan | Onion | Oshima et al., 2004 |
| OY-V | GCF_000744065 | Illumina and Roche/454; 300x | 170 contigs | 739,592 | 651 | 217 | Japan | Onion | Kakizawa et al., 2014 |
| DY2014 | GCA_005093185 | Illumina; 124x | 8 contigs | 824,596 | 775 | 63 | Taiwan | Periwinkle | Cho et al., 2019b |
| MBSP-M3 | GCF_001712875 | Illumina; 42x | Complete | 576,118 | 498 | 34 | Brazil | Maize | Orlovskis et al., 2017 |
| De Villa | GCF_004214875 | Illumina; 50x | Complete | 603,949 | 512 | 21 | South Africa | Grapevine | Coetzee et al., 2019 |
| LD1 | GCF_001866375 | Illumina; 200x | 8 contigs | 599,264 | 536 | 29 | China | Rice | Zhu et al., 2017 |
| CYP | GCF_000803325 | Illumina; 47x | 252 contigs | 659,699 | 546 | 154 | Italy | Daisy | Pacifico et al., 2015 |
| TW1 | GCA_003181115 | Illumina and ONT; 146x | 5 contigs | 743,598 | 457 | 248 | Canada | Canola | Town et al., 2018 |

## Materials and Methods

### Data Sets and Analysis Methods

The 11 genome sequences included in this study are listed in Table 1. These data sets represent all phytoplasma genomes available from GenBank (Benson et al., 2018) as of January 2020 and are recognized as belonging to the 16SrI group, AY group, or ‘*Ca*. P. asteris’ (Lee et al., 2004). Two other ‘*Ca*. Phytoplasma’ species have been described as being affiliated with the 16SrI group, including ‘*Ca*. P. japonicum’ (Sawayanagi et al., 1999) and ‘*Ca*. P. lycopersici’ (Arocha et al., 2007), but were not included in this study due to the lack of genome sequence availability. Notably, ‘*Ca*. P. japonicum’ was first reported as a member of the 16SrI-D subgroup (Sawayanagi et al., 1999) and later listed under the 16SrI group in the current taxonomy (The IRPCM Phytoplasma/Spiroplasma Working Team, 2004). However, the *i*PhyClassifier (Zhao et al., 2009) analysis of its 16S rRNA gene sequence (GenBank accession AB010425) assigned it to the 16SrXII-D subgroup, suggesting that its classification remained to be investigated.

To confirm the 16Sr group and subgroup assignments, we extracted the 16S rRNA gene sequences from the genomes and analyzed those sequences using the *i*PhyClassifier (Zhao et al., 2009). Most phytoplasma genomes contain two copies of 16S rRNA genes and each copy was processed individually. The strain OY-V lacks any 16S rRNA gene in the current assembly and therefore was excluded from this analysis. The strain NJAY has four partial sequences for its 16S rRNA genes and all of these four sequences were examined.

All bioinformatic tools for processing these data sets were used with the default settings unless stated otherwise. For correlation tests, the cor.test function implemented in R v3.4.4 (R Core Team, 2019) was used to examine the Pearson’s product moment correlation coefficient.

For strain TW1 (GenBank genome accession GCA_003181115.1), we downloaded the raw sequencing results from the NCBI Sequence Read Archive (SRA) to examine the phytoplasma sequences present in the sequencing libraries. The raw reads were mapped to two other reference genomes (i.e., AYWB and OY-M; see Table 1) to detect the presence of 16SrI-A and 16SrI-B type phytoplasmas, respectively. For the Oxford Nanopore MinION reads (SRA accession SRR7548026), the mapping was performed using Minimap2 v2.15 (Li, 2018). For the Illumina MiSeq paired-end reads (SRA accession SRR7548027), the mapping was performed using the Burrows-Wheeler Alignment (BWA) tool v0.7.17 (Li and Durbin, 2009). The mapping results were programmatically checked using SAMtools v1.9 (Li et al., 2009) to identify the sequence variations.

### Genome Comparisons

For whole-genome comparison, FastANI v1.1 (Jain et al., 2018) was used to calculate the proportion of genomic segments mapped and the average nucleotide identity (ANI) of those segments in each genome-pair. For each pair, reciprocal comparisons between the two genomes were conducted (i.e., X as the query against Y as the reference and Y as the query against X as the reference). To calculate the proportion of genomic segments mapped, the numbers of segments mapped from those two reciprocal comparisons were combined as the numerator, and the total numbers of segments from those two genomes were combined as the denominator. The reciprocal ANI values were averaged. For example, in the AYWB-NJAY comparison, 183 out of the 239 segments of AYWB could be mapped to NJAY, and all 187 segments of NJAY could be mapped to AYWB. Based on these results, these two strains share (183 + 187) / (239 + 187) = 370 / 426 = 86.9% of their genomic segments. The ANI values were 99.5% and 99.8% in the two reciprocal comparisons, and an average ANI of 99.7% was reported.

For gene-centric investigation, the homologous gene clusters among all genomes were identified using OrthoMCL v1.3 (Li et al., 2003) based on the procedures described in our previous studies (Lo et al., 2013; Chung et al., 2013). Briefly, all coding sequences (CDS), including those annotated as pseudogenes, were included in the all-against-all similarity searches using BLASTN v2.6.0 (Camacho et al., 2009) with an e-value cutoff of 1e^-15^. The BLASTN results were used as the input for OrthoMCL, which normalized the within- and between-genome comparisons and identified homologs according to the principle of reciprocal best hits. Similar to the procedures for whole-genome comparisons, the proportion of homologs shared and the sequence identity of shared homologs were considered as two separate metrics of genome similarities. The proportion of homologs shared was calculated based on averaging (i.e., the number of homologs shared was used as the numerator, and the average number of homologs present in those two genomes was used as the denominator). The sequence identities of coding regions were calculated based on multiple sequence alignment of those single-copy genes present in all of the genomes compared. The homologs were aligned using MUSCLE (Edgar, 2004) and the sequence identities were calculated using the DNADIST program in PHYLIP v3.697 (Felsenstein, 1989); as expected, reciprocal comparisons all produced the same result for sequence identities.

In addition to the pairwise comparisons, the gene content of all genomes was compared using a procedure based on principal coordinates analysis (Lo et al., 2018). For this procedure, the homologous gene clustering result was converted into a matrix of 11 genomes by 1,233 gene clusters; the value in each cell indicates the gene copy number. This matrix was converted into a Jaccard distance matrix among genomes using the VEGAN package in R, then processed using the PCOA function in the APE package (Popescu et al., 2012).

To compare the genes for putative secreted proteins among these phytoplasmas, we examined all coding sequences using a procedure described in a previous study (Cho et al., 2019b). Briefly, SignalP v5.0 (Armenteros et al., 2019) was used to check the presence of signal peptide based on the Gram-positive bacteria model. The program TMHMM v2.0 (Krogh et al., 2001) was used to verify that these putative secreted proteins have no transmembrane domain. For those four experimentally characterized effectors (i.e., SAP05, SAP11/SWP1, SAP54/PHYL1, and TENGU), all putative homologs (i.e., those belong to the same homologous gene cluster as defined by OrthoMCL or have a BLASTN e-value of lower than 1e^-100^) were manually added to the list of regardless of the SignalP prediction results.

The genome alignment of representative strains with high quality assemblies were performed using genoPlotR v0.8.9 (Guy et al., 2010). The syntenic regions were identified using BLASTN v2.6.0 (Camacho et al., 2009), the cutoff values for sequence similarity were set to e-value = 1e^-15^ and alignment length = 2,000 bp.

### Molecular Phylogenetics

For phylogenetic inference, MUSCLE v3.8.31 (Edgar, 2004) was used to generate multiple sequence alignments and PhyML v3.3 (Guindon and Gascuel, 2003) was used to infer maximum likelihood phylogenies. The visualization was performed using JalView v2.11 (Waterhouse et al., 2009) and FigTree v1.4.4. In each inference, PHYLIP v3.697 (Felsenstein, 1989) was used to generate 1,000 replicates for bootstrap analysis.

### Development of Molecular Markers

To identify molecular markers that may be suitable for classification, those multiple sequence alignments of single-copy genes shared by all genomes were examined manually. Genes were ranked by the number of variable sites that could be used for distinguishing different OTUs. Subsequently, the list was prioritized based on the criterion of having multiple markers located in distinct regions of those phytoplasma chromosomes. The PCR primers (Table 2) were designed manually with the aim of producing ∼700-800 bp products to cover those variable regions, such that each product could be sequenced in one single Sanger sequencing reaction.

**Table 2.**
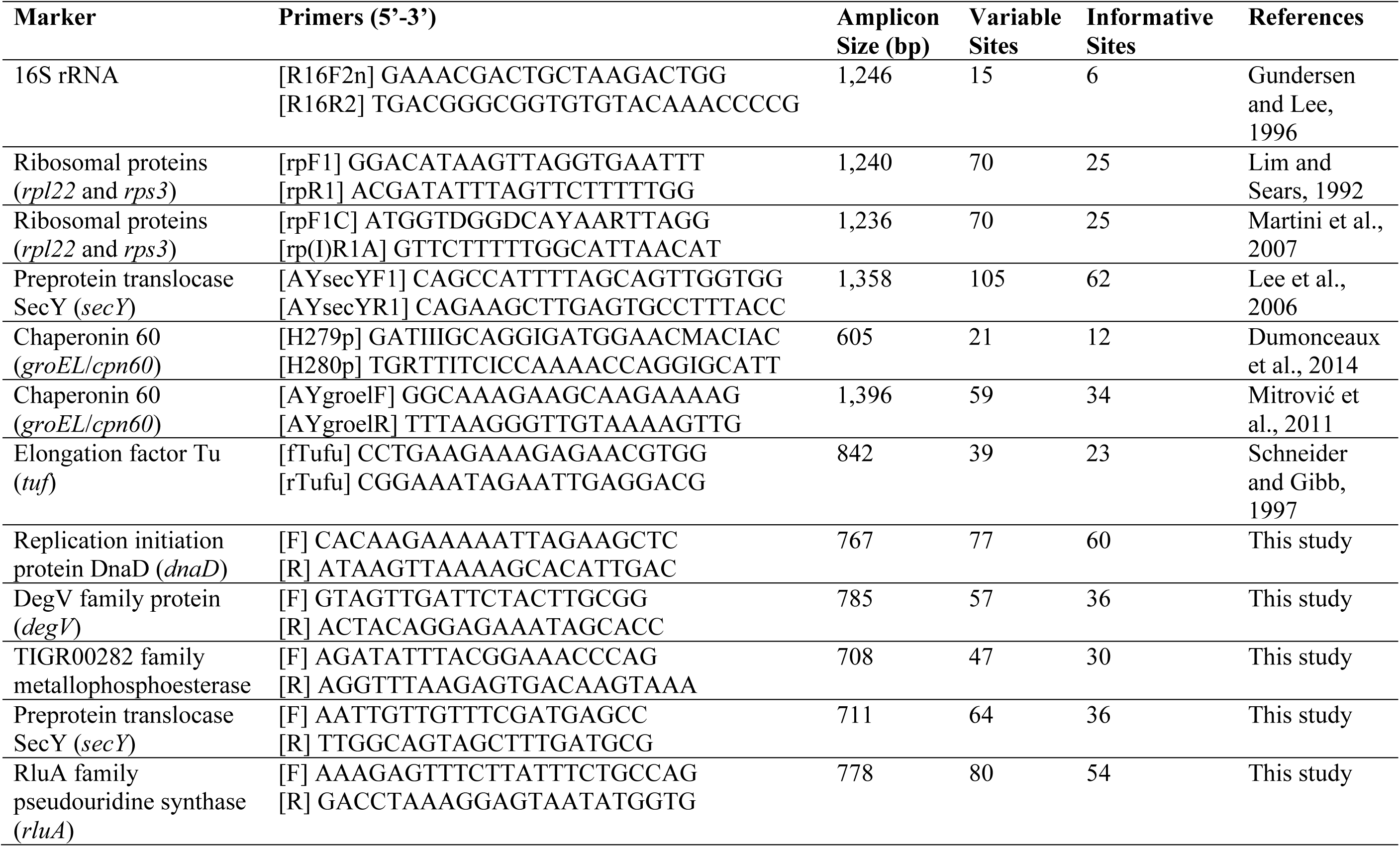
List of molecular markers for phytoplasma classification.

To validate the primer design, a DNA sample derived from a periwinkle infected by the phytoplasma strain DY2014 (Cho et al., 2019b) was used as the template for PCR tests. All kits were used according to the manufacturer’s instructions. The PCR were performed using the KAPA HiFi HotStart ReadyMix (Roche, USA). The cycling conditions were: (1) an initial activation step at 95°C for 3 min; (2) 35 cycles of 98°C for 20 sec, 55°C for 30 sec, and 72°C for 60 sec; and (3) a final extension step of 72°C for 5 min. The annealing temperature was lowered to 53°C for the newly designed primers for *secY* (Table 2), while the other four primer sets used 55°C. Agarose gel electrophoresis was conducted to confirm that all PCR tests produced one single band of expected size. For further confirmation, the PCR products were purified using the QIAquick Gel Extraction Kit (Qiagen, Germany), then sequenced using the BigDye Terminator v3.1 Cycle Sequencing Kit on an Applied Biosystems 3730XL DNA Analyzer (Thermo Fisher Scientific, USA). The sequencing results were compared to the published genome sequence for final confirmation.

The procedure for calculating the substitution rates was based on that described in our previous studies (Kuo and Ochman, 2009; Kuo et al., 2009). Briefly, the homologous protein sequences were aligned by using MUSCLE v3.8.31 (Edgar, 2004). The protein alignments were converted into codon-based nucleotide alignment using PAL2NAL v14 (Suyama et al., 2006), which were then processed using the YN00 method (Yang and Nielsen, 2000) implemented in PAML v4.9h (Yang, 2007). To identify the homologs of selected marker genes in other more divergent phytoplasmas, the full gene sequences were used as queries to run BLASTN searches against available genomes. After the homolog identification, multiple sequence alignments were performed using MUSCLE (Edgar, 2004) and the newly designed primers (Table 2) were examined manually to check if these primers could work on other phytoplasmas.

## Results and Discussion

### Genome Characteristics

Among the 11 phytoplasma genomes included in this study, four have the complete chromosomal sequence available while the other seven are draft assemblies with varying levels of completeness (Table 1 and Figure 1). For those incomplete genomes, the true genome sizes are difficult to determine. Nonetheless, the largest (i.e., OY-M; 853 kb) and the smallest (i.e., MBSP-M3; 576 kb) genomes in this data set are both completed assemblies, confirming the high level of genome size variation among these closely related strains. The total number of CDS is strongly correlated with the genome size (r = 0.92, p = 2.7e-5). However, the number and proportion of pseudogenes varied widely among these genomes. For example, the strains AYWB and NJAY have a genome-wide ANI of 99.7% (Figure 2A), yet the proportion of pseudogenes are 17% and 41% (Table 1), respectively. In another case, DY2014 and OY-V have a genome-wide ANI of 99.4%, yet the proportion of pseudogenes are 8% and 25%, respectively. Although these differences may reflect true biological processes, such as elevated mutation accumulations that have occurred in the evolutionary history of some strains, it is also possible that these differences may be explained by artifacts. Regarding the sequencing methods, AYWB was based on Sanger sequencing (Bai et al., 2006) and DY2014 was based on Illumina (Cho et al., 2019b), while NJAY was based on one single Roche/454 library (Sparks et al., 2018) and OY-V was based on one Illumina paired-end library and one Roche/454 GS FLX mate-pair library (Kakizawa et al., 2014). Due to the low GC-content of these genomes (i.e., ∼27-28%), it is plausible that sequencing errors in homopolymeric regions, particularly those errors originated from the 454 sequencing technology, may have contributed to the high numbers of pseudogenes with frameshift mutations in NJAY and OY-V. Moreover, assemblies that are more fragmented may also have more partial genes located at the ends of contigs. Due to these concerns, all pseudogenes were included in the gene-centric investigation of this study.

**Figure 1.**
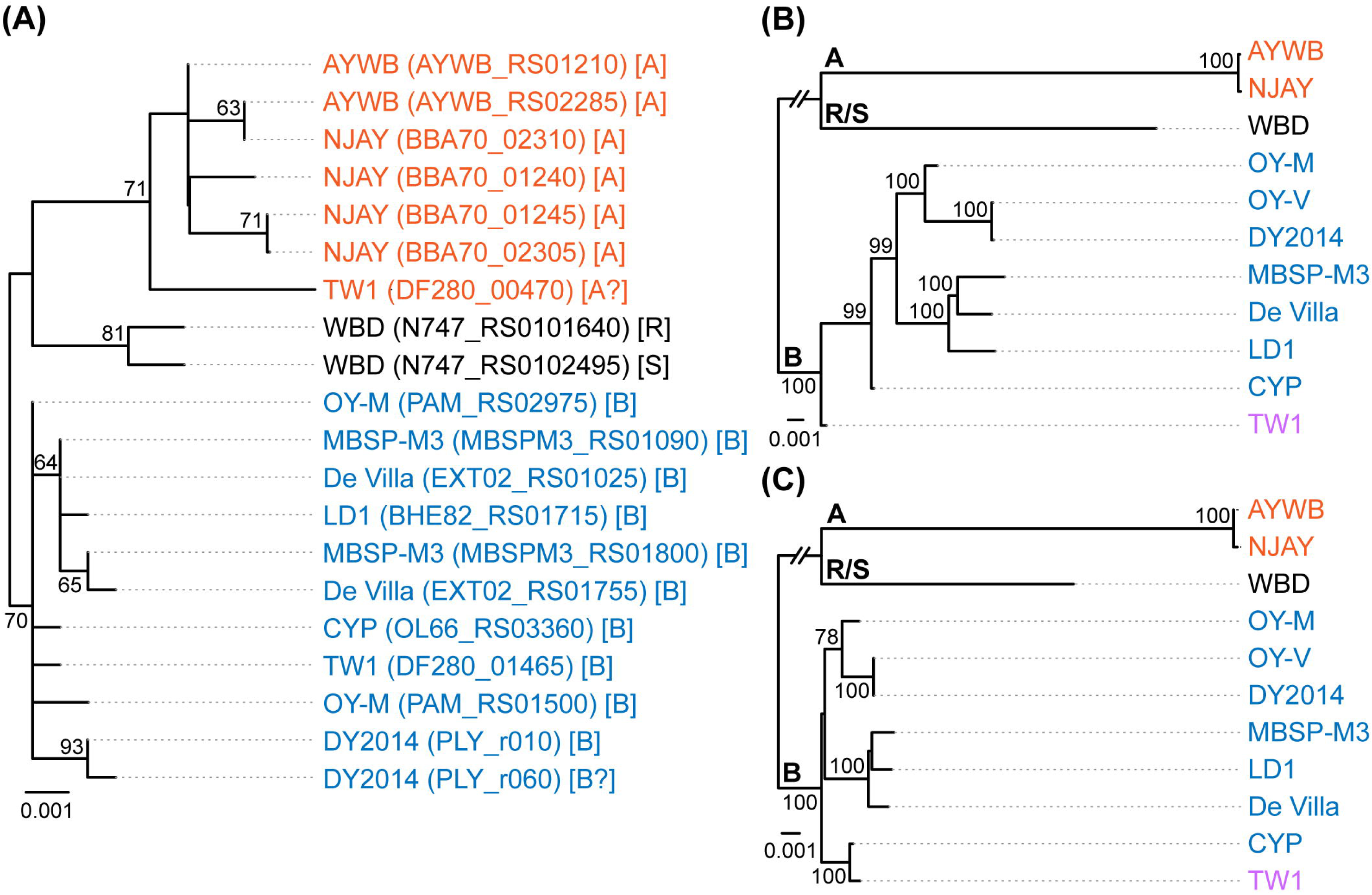
Maximum likelihood phylogenies. The numbers on the internal branches indicate the level of bootstrap support based on 1,000 resampling; only values >=60% are shown. (A) 16S rRNA genes. The alignment contains 1,541 aligned nucleotide sites. The strain name, locus tag of the 16S rRNA gene (in parentheses), and the 16SrI subgroup classification (in square brackets) are labeled. A question mark ‘?’ in the subgroup classification indicates that the sequence is classified as a new subgroup, the existing subgroup with the highest similarity is provided. (B) Single-copy coding genes shared by all strains. The concatenated alignment contains 303 genes and 291,990 aligned nucleotide sites. (C) 16S rRNA gene and the five markers developed in this study; see Table 2 for detailed information. The concatenated alignment contains 4,995 aligned nucleotide sites.

**Figure 2.**
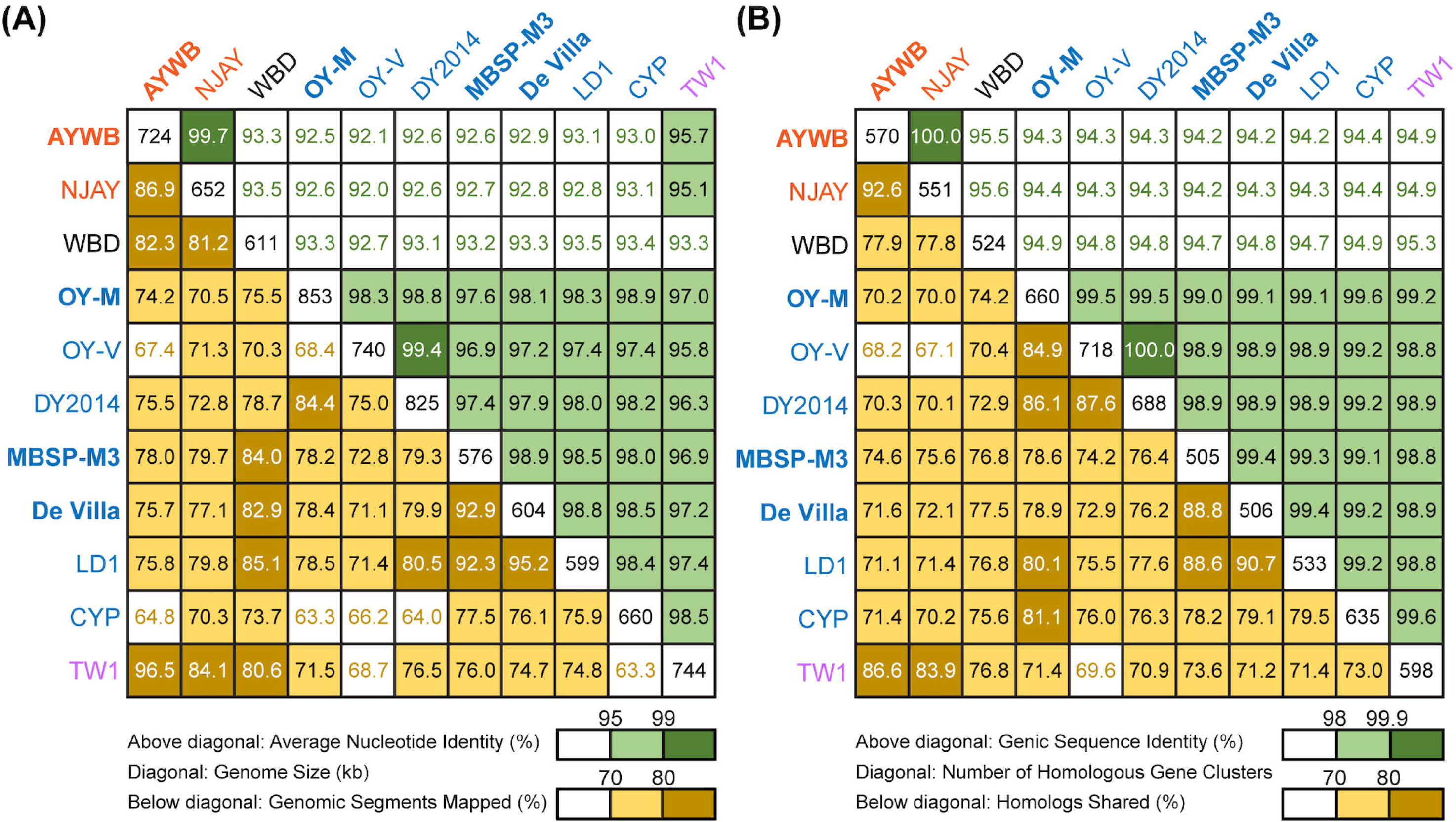
Pairwise genome similarities. The strains with the complete genome sequences available are highlighted in bold. (A) Similarity scores based on whole genome comparisons. Above diagonal: average nucleotide identity (%); diagonal: genome size (kb); below diagonal: genomic segments mapped (%). (B) Similarity scores based on genic regions. Above diagonal: genic sequence identity (%); diagonal: number of homologous gene clusters; below diagonal: homologs shared (%).

### Classification of the 16SrI Phytoplasmas Analyzed

The 16SrI subgroup assignments of these strains were mostly consistent with those reported in literature but there were a few surprises (Figure 1A). First, WBD was reported as a 16SrI-B strain (Chen et al., 2014). However, our examination of the sequence record deposited in GenBank revealed that the two copies of 16S rRNA genes in this genome differ by 4-bp, and *i*PhyClassifier (Zhao et al., 2009) assigned these two sequences to 16SrI-R and 16SrI-S, respectively. Results from molecular phylogeny also support that these two sequences form a clade that is distinct from other 16SrI-B sequences. Second, DY2014 has two 16S rRNA genes that differ by 1-bp. While one was assigned to 16SrI-B, the other was identified as being a new subgroup with a RFLP pattern most similar to 16SrI-B. Finally, TW1 has two sequences that differ by 12-bp, with one assigned to 16SrI-A and the other assigned to 16SrI-B (Town et al., 2018).

These results highlighted several issues of this RFLP-based classification system for phytoplasmas. First, the intra-genomic variation of 16S rRNA genes may result in a strain being assigned to different subgroups depending on which homolog was examined. This issue has been reported for other phytoplasmas (Liefting et al., 1996) and a three-letter subgroup designation system has been proposed (Wei et al., 2008). For example, a paulownia witches’-broom (PaWB) phytoplasma was first classified as a 16SrI-D strain (Lee et al., 1998) but later found to harbor a 16SrI-B type sequence as well. To accommodate these findings, the subgroup status of this PaWB was redesignated as 16SrI-(B/D)D, with the first two letters in parentheses denoting the types of those two 16S rRNA genes, and the third letter denoting its 16Sr subgroup assignment (Wei et al., 2008; Zhao et al., 2009). Although this three-letter system provides comprehensive information, it may be unnecessarily complex and confusing. Moreover, it is unclear which one of the two types should be chosen for the formal subgroup assignment. In the case of PaWB, it was assigned to 16SrI-D based on precedence. However, in the case of WBD (i.e., originally described as a 16SrI-B strain), it is unclear if it should be redesignated as 16SrI-(R/S)R or 16SrI- (R/S)S. Second, this RFLP-based system considers only the sequence variations in restriction sites. By ignoring the sequence variations in other positions, this approach could alleviate the problem of intra-genomic variations to a certain degree (e.g., the two sequences in AYWB differ by 2-bp, yet both were assigned to 16SrI-A). However, this approach wasted much of the information contained in the full sequence and the uneven weighing of nucleotide positions could introduce strong biases. The strain DY2014 represents one extreme example to illustrate such biases, with a single-bp difference that is sufficient to result in different subgroup assignments between the two 16S rRNA genes. Finally, a classification system that depends solely on the 16S rRNA genes while ignoring other regions of the genome does not provide sufficient resolution.

Regardless of the group/subgroup assignments of these 16S rRNA genes, all these sequences share >99% identity (Supplementary Table S1), which is above the general recommended thresholds for defining species. These thresholds include: (1) 97% identity that is commonly used for defining OTUs corresponding to species in microbiota surveys (Schloss and Handelsman, 2005), (2) 97.5% identity recommended for ‘*Ca*. Phytoplasma’ species (The IRPCM Phytoplasma/Spiroplasma Working Team, 2004), and (3) 98.6% identity recommended for uncultivated bacteria (Konstantinidis et al., 2017). In other words, these strains may be considered as all belonging to ‘*Ca*. P. asteris’ (Lee et al., 2004) based on their 16S rRNA gene sequences. However, delineating species-level OTUs based on one single gene may be problematic; more in-depth investigation of their genomic divergence is necessary.

### Genomic Divergence among 16SrI Phytoplasmas and Putative Species Boundaries

To further investigate the divergence of these phytoplasmas, we utilized two approaches to quantify their genomic similarities. In the first approach, the whole genome sequences were used to calculate the proportion of genomic segments shared, as well as the ANI values of these segments. A previous study that examined >90,000 prokaryotic genomes found that the ANI values exhibit a bimodal distribution, and the within-species comparisons almost always have >95% ANI (Jain et al., 2018). We found that based on this criterion, the 11 strains examined could be separated into three clusters that roughly correspond to their 16SrI subgroup assignments (Figure 2A). The two 16SrI-A strains, AYWB and NJAY, share 86.9% of their genomes and these shared regions are nearly identical (i.e., ANI = 99.7%). Because the NJAY assembly is highly fragmented and ∼10% smaller compared to the complete genome of AYWB, the low proportion of genomic segments shared may be explained by missing data in the NJAY assembly. The strain WBD shares only ∼93% ANI with all other 16SrI phytoplasmas compared and represents a distinct lineage. All of the remaining strains, including those assigned to 16SrI-B and TW1, share >97% ANI and could considered as a third cluster. This clustering result is consistent with the maximum likelihood phylogeny inferred using 303 single-copy genes shared by all strains (Figure 1B and Supplementary Table S2).

In the second approach, we considered only the putative coding regions (i.e., intact CDS and putative pseudogenes) of these genomes for calculating genomic similarities (Figure 2B). Compared to the genome-wide ANI approach, this gene-centric method is more laborious and could potentially be affected by annotation quality. Nevertheless, this approach has a higher confidence in inferring the homology prior to calculating sequence identities and provides a complementary method to the ANI approach. Although the exact genome similarities values differ slightly, the patterns found by using these two approaches are consistent.

Based on these results regarding genomic similarities and molecular phylogeny, it appeared that these 16SrI phytoplasmas could be classified into three species-level OTUs. These findings are consistent with a previous study that the conventional 97% identity threshold for delineating species in bacteria based on 16S rRNA gene is too low (Edgar, 2018). Similar to a previous study that examined diverse prokaryotes (Jain et al., 2018), the sequence identity values among these phytoplasmas also exhibited a bimodal distribution (Figure 3). Excluding the comparisons involving TW1, the within-OTU comparisons all have >97% genome-wide ANI, while between-OTU comparisons all have <94% ANI (Figures 2A and 3A). For sequence identities that were calculated based on only the genic regions, in which the homology could be inferred more confidently, a similar separation was found (i.e., within-OTU: >98.9%, between-OTU: <96%; see Figures 2B and 3B). These results suggest that those strains assigned to the same species-level OTU may have on-going homologous recombination that maintained high sequence similarities in their shared genomic regions. In contrast, certain genetic barriers exist to lower homologous recombination between different OTUs. In other words, these results provide further support to the hypothesis that the biological species concept, which is based on the barriers to homologous gene exchange, could be applied to bacteria as well (Bobay and Ochman, 2017). Alternatively, other hypotheses that may explain the discontinuity in genome similarities include ecological sweeps or stochastic neutral processes (Jain et al., 2018), and the exact biological mechanisms remained to be investigated.

**Figure 3.**
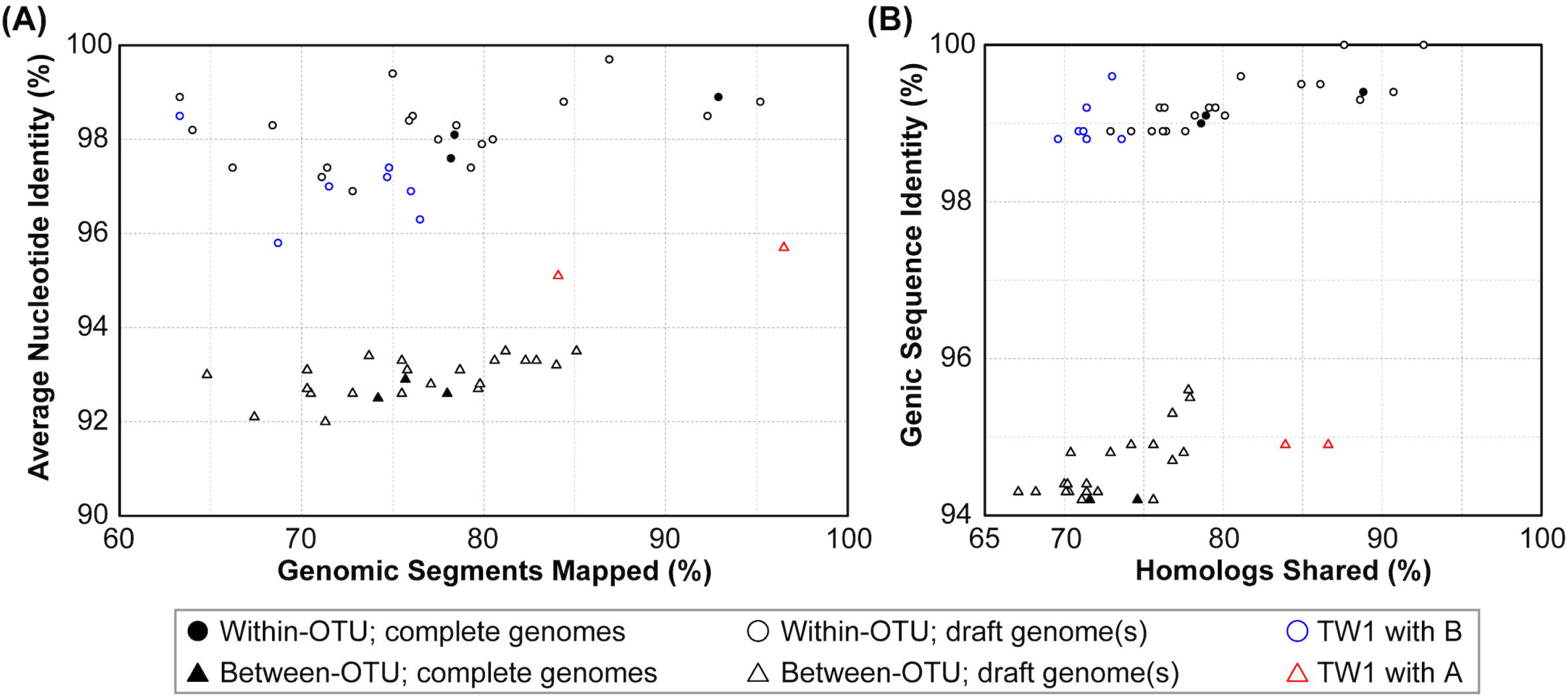
Correlations between metrics of genome similarities. (A) Similarity scores based on whole genome comparisons. (B) Similarity scores based on genic regions. Filled symbols indicate the pairwise comparisons involving two complete genomes, empty symbols indicate the pairwise comparisons involving one or two draft genomes. The strain TW1 is a special case; its similarity scores in comparison with 16SrI-A and 16SrI-B strains are indicated by red and blue, respectively.

In addition to sequence identity, another important measurement for genetic similarity is the proportion of genomic regions shared. For this second measurement, a wide range was observed for both within- and between-OTU comparisons (Figures 2 and 3). The wide spread was mostly explained by the inclusion of two highly fragmented draft genomes (i.e., OY-V and CYP; see Table 1), which resulted in underestimates that are artifacts stemming from missing data. However, because the non-redundant chromosomal regions that are more conserved have a higher probability of being included in a draft assembly, the inclusion of draft genomes may also result in overestimates of the genomic regions shared. Due to these uncertainties, we restricted the comparisons to those four strains with the complete genome sequences available (i.e., AYWB, OY-M, MBSP-M3, and De Villa). For this reduced data set with high confidence, strains assigned to the same species-level OTU share ∼78-93% of their chromosomal segments and ∼79-89% of their protein-coding genes, whereas strains assigned to different OTUs share ∼74-78% of their chromosomal segments and ∼70-75% of their protein-coding genes.

For an alternative approach of comparing the gene content divergence among these phytoplasmas, we conducted a principal coordinates analysis that considered gene copy numbers in addition to the patterns of gene presence or absence (Figure 4). With the exception of TW1, the results were consistent with the molecular phylogeny (Figure 1B) and pairwise genome similarities (Figures 2 and 3). The first coordinate explained ∼40% of the variance and showed clear separation for the three species-level OTUs. The second coordinate explained ∼23% of the variance and showed that those 16SrI-B strains could be separated into three subgroups.

**Figure 4.**
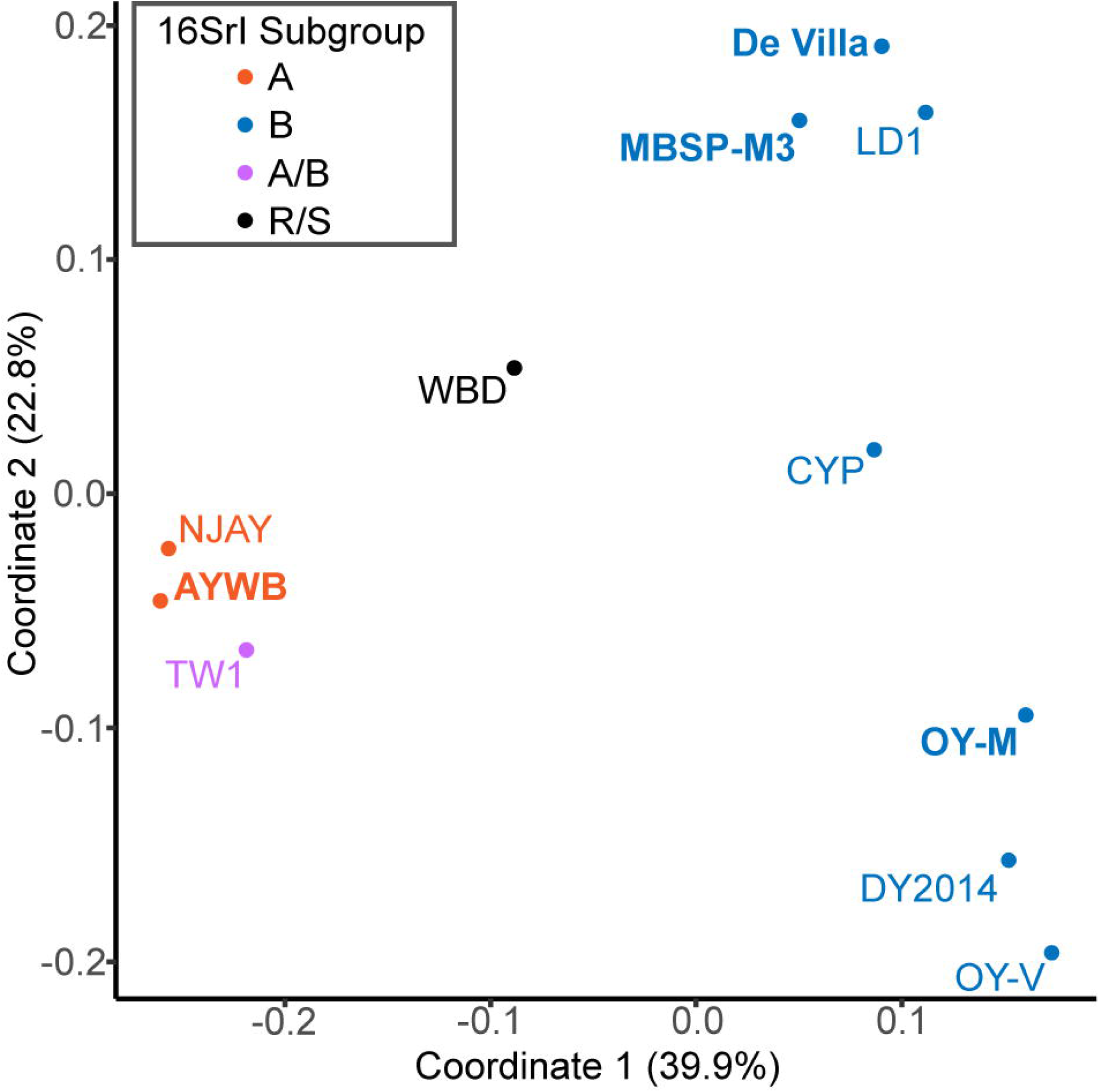
Principal coordinates analysis of gene content. Each genome is represented by one dot (color-coded according to the 16Sr subgroup assignment). The distance between dots corresponds to their differences in gene content. The % variance explained by each axis is provided in parentheses. The strains with the complete genome sequences available are highlighted in bold.

Taken together, these results supported the existence of multiple species-level OTUs within the previously described ‘*Ca*. P. asteris’ (Lee et al., 2004) and provided quantitative guidelines for inferring the putative species boundaries based on overall genome similarities. Based on these findings, the critical questions to ask are the genotypic, phenotypic, and ecological differentiations among these OTUs. Unfortunately, given the limited number of strains with genome sequence available, it is not possible to obtain meaningful answers yet. For example, other 16SrI-A strains were known to infect different dicots or monocots, and were found in North America and Europe (Lee et al., 2004). However, no genomic information is available for these strains with different hosts and/or geographic distributions. It is important to characterize these strains and validate their OTU assignment before meaningful comparisons could be made between different OTUs. Similarly, more strains that belong to the same OTU with WBD must be sampled and characterized before the biological characteristics of this OTU could be described appropriately. These improvements in the understanding of genotypic and phenotypic differentiations among phytoplasmas are critical in informing future taxonomy revisions.

### The Special Case of the TW1 Genome

Based on the aforementioned framework for phytoplasma classification, the strain TW1 represents a strange and difficult case. The high level of intra-genomic variation between its two copies of 16S rRNA genes has been noted in the initial characterization (Town et al., 2018). In our whole-genome analysis, this strain shares a higher proportion of its chromosomal segments and protein-coding genes with those 16SrI-A strains, yet has higher sequence identities with those 16SrI-B strains (Figures 2 and 3). Because of these properties, this strain was clustered with those 16SrI-A strains based on gene content (Figure 4), while having closer phylogenetic relationships with those 16SrI-B strains (Figure 1B).

In the absence of an intuitive hypothesis that could explain the evolutionary processes leading to these conflicting findings, we examined the possibilities that these findings were results of artifacts. This TW1 genome was co-assembled using one long-read library (based on Oxford Nanopore MinION) and one short-read library (based on Illumina MiSeq) (Town et al., 2018). When the raw reads from these two sequencing libraries were examined separately, we found that the long-read library likely contains a 16SrI-A type phytoplasma. When the raw reads were mapped to the AYWB chromosome, all regions were covered and 3,725 sequence polymorphisms were found; when mapped to OY-M, 52,649 bp had no coverage and 31,401 sequence polymorphisms were found. In contrast, the short-read library likely contains a 16SrI-B type phytoplasma. When the raw reads were mapped to OY-M, 103,922 bp had no coverage and 9,489 sequence polymorphisms were found; when mapped to AYWB, 110,458 had no coverage and 29,323 sequence polymorphisms were found.

Based on these findings, a possible explanation is that during the assembly process, the long reads produced several 16SrI-A type scaffolds, which resulted in the high similarities of the TW1 genome content to 16SrI-A as observed (Figures 2-4). Subsequently, during the polishing stage, the exact sequences of these scaffolds were modified based on the short-read library, thus explaining the high sequence similarity of this TW1 genome to those 16SrI-B strains. In other words, in its current form, this TW1 genome assembly is likely an artifact that was incorrectly assembled by combining two sequencing libraries with phytoplasmas belonging to different OTUs, rather than a true representative of an existing phytoplasma strain. Due to the concern that the inclusion of this genome may have introduced biases, we repeated all phylogenetic inferences and excluded the TW1 sequences prior to multiple sequence alignment. The resulting tree topologies were consistent in terms of OTU assignments; the minor differences were related to the placement of CYP (Figure 1 and Supplementary Figure 1).

### Development of Molecular Markers for Multilocus Sequence Analysis

To develop molecular markers that may be useful for the MLSA of these phytoplasmas and their relatives, we examined the multiple sequence alignments of those 303 single-copy genes shared by all of the genomes analyzed. The selection criteria were based on: (1) the number of informative sites that may be used to distinguish the three species-level OTUs identified, (2) the possibility of designing PCR primers for a product in the size range of ∼700-800 bp and covers a large number of informative sites, and (3) the chromosomal locations. The first two criteria were aimed to make the Sanger sequencing of these markers as cost-effective as possible, while the third criterion was aimed to make the MLSA robust against recombination.

Based on these criteria, we selected five markers to supplement the 16S rRNA gene. These five makers all have high densities of informative sites, particularly when compared to those markers that have been developed previously (Table 2 and Supplementary Figure S1). The substitute rates of these marker genes are relatively high but do not deviate strongly when compared to other shared single-copy genes (Supplementary Figure S2).

Based on the functional annotation, two of these selected markers (i.e., DegV family protein and TIGR00282 family metallophosphoesterase) are not generally considered as house-keeping genes. These results reflected our philosophy of marker selection (i.e., prioritize the candidates with the highest resolving power for the target taxa regardless of other irrelevant attributes). Similar to our criteria and results, a pioneer study on genome-enable MLSA marker design in phytoplasma selected several genes that were annotated as hypothetical proteins (Kakizawa and Kamagata, 2014). In terms of phylogenetic distribution, we found that four of these five selected genes are present as single-copy genes in other phytoplasmas, including ‘*Ca*. P. australiense’ of the 16SrXII group (Tran-Nguyen et al., 2008), ‘*Ca*. P. aurantifolia’ of the 16SrII group (Chung et al., 2013), ‘*Ca*. P. pruni’ of the 16SrIII group (Lee et al., 2015), ‘*Ca*. P. ziziphi’ of the 16SrV group (Wang et al., 2018a), ‘*Ca*. P. oryzae’ of the 16SrXI group (GenBank accession NZ_JHUK00000000), and ‘*Ca*. P. mali’ of the 16SrX group (Kube et al., 2008). The only exception is *degV*, which is absent in two of the draft genomes (i.e., ‘*Ca*. P. aurantifolia’ and ‘*Ca*. P. oryzae’; may be located in the unassembled regions) and has two tandem copies in the complete genome sequence of ‘*Ca*. P. mali’. Nevertheless, the sequence divergence levels are too high for all these genes, such that the primers designed in this study (Table 2) may not work for these more divergent phytoplasmas outside of the 16SrI group. These results are consistent with our expectations that different MLSA markers are necessary for different phylogenetic depths.

In terms of chromosomal locations, these markers are well separated in all genomes examined despite the extensive genome rearrangements observed (Figure 5). Although the extent of recombination in phytoplasmas is not well understood, recent studies on *Xylella fastidiosa* (i.e., another insect-transmitted plant pathogen that is restricted to vascular tissues) revealed that recombination affected ∼0.5-15% of the genome, depending on the exact subspecies (Potnis et al., 2019; Vanhove et al., 2019). The recombined fragments could have sizes up to 31 kb, with an average of 1 kb. Based on these estimates, it is unlikely that one single recombination event would affect more than one of these five markers. Thus, the results of MLSA should be robust against the interference of recombination.

**Figure 5.**
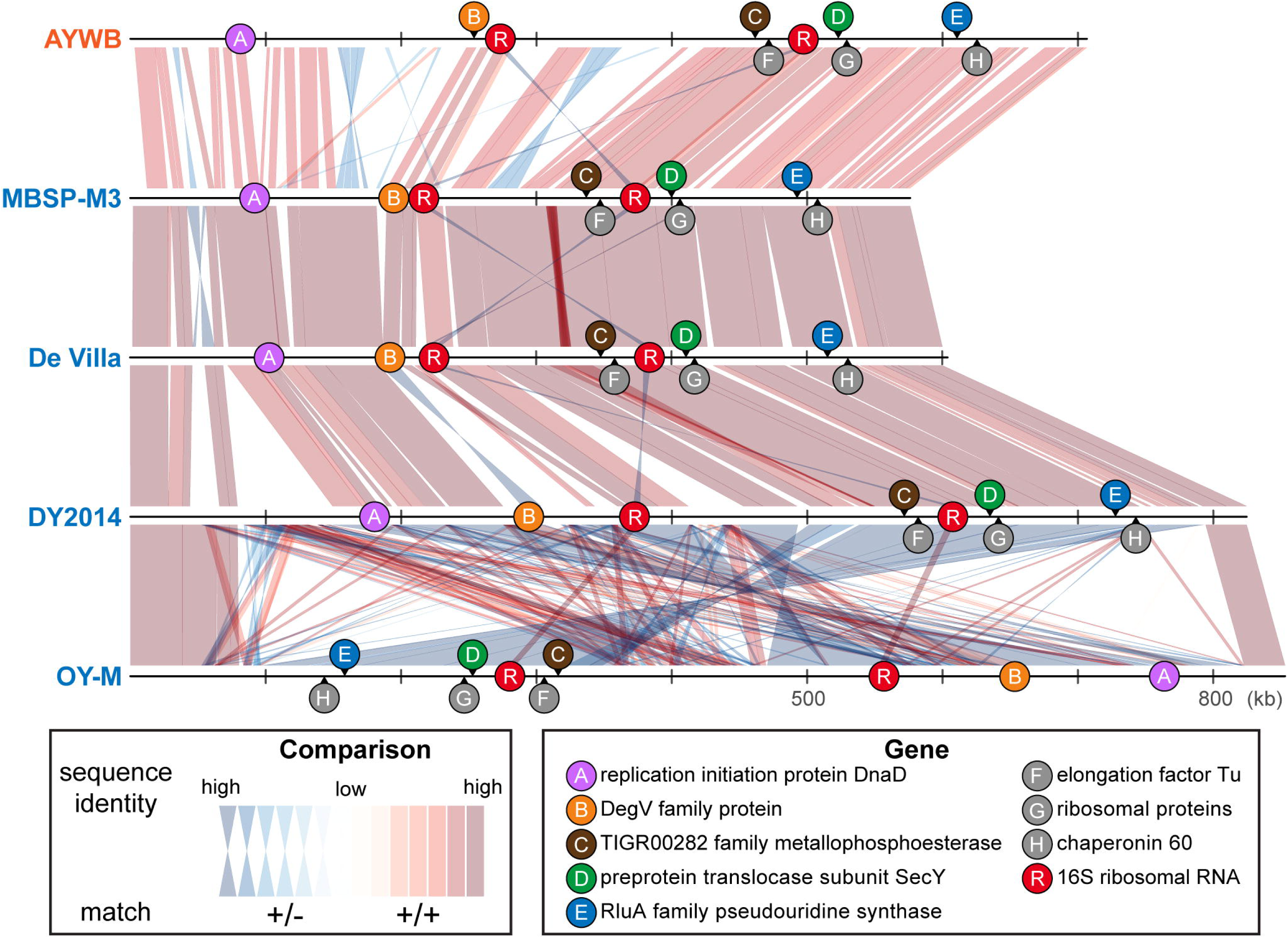
Genomic locations of marker genes. Nucleotide sequence identity of the conserved regions between genomes are illustrated (red: match on the same strand; blue: match on the opposite strand).

When we used these five newly developed markers for molecular phylogenetic inference, the concatenated alignment (together with the 16S rRNA gene, which is expected to be sequenced in the initial characterization of newly collected strains) produced a phylogeny (Figure 1C and Supplementary Figure 1C) that is comparable to the result from genome-scale analysis (Figure 1B and Supplementary Figure 1B). This result demonstrated the usefulness of these markers for future genotyping work. With better sampling of phytoplasma strains and more accurate classification of those strains using these markers, informed decision could be made to select representatives for genome sequencing efforts, which could contribute to the study of phytoplasma genetic diversity and evolutionary history, as well as improve the taxonomy. Moreover, as the sampling focus changes in the future (e.g., a higher resolution is desired for one of the OTUs), the list of conserved single-copy genes and the substitution rates of those genes (Supplementary Table S2) could provide a guide for developing more suitable MLSA markers.

### Phylogenetic Distribution of Phytoplasma Effector Genes

With the well-resolved phylogeny (Figure 1B) as a framework, we investigated the distribution of putative effector genes among these phytoplasmas. The results indicated a high level of variation in the gene counts (Figure 6 and Supplementary Table S3). One clade (i.e., MBSP-M3, De Villa, and LD1) have only ∼10-14 genes that encode putative secreted proteins. These numbers are much lower than the ∼36-45 found in their sister clade (i.e., OY-M, OY-V, and DY2014). The low numbers are not artifacts of incomplete genome sequences. In fact, the two strains with the lowest numbers (i.e., MBSP-M3 and De Villa) both have the complete chromosomal sequences available. These findings suggest that there may have been lineage-specific reduction or expansion in effector gene counts, although the underlying ecological factors are unclear.

**Figure 6.**
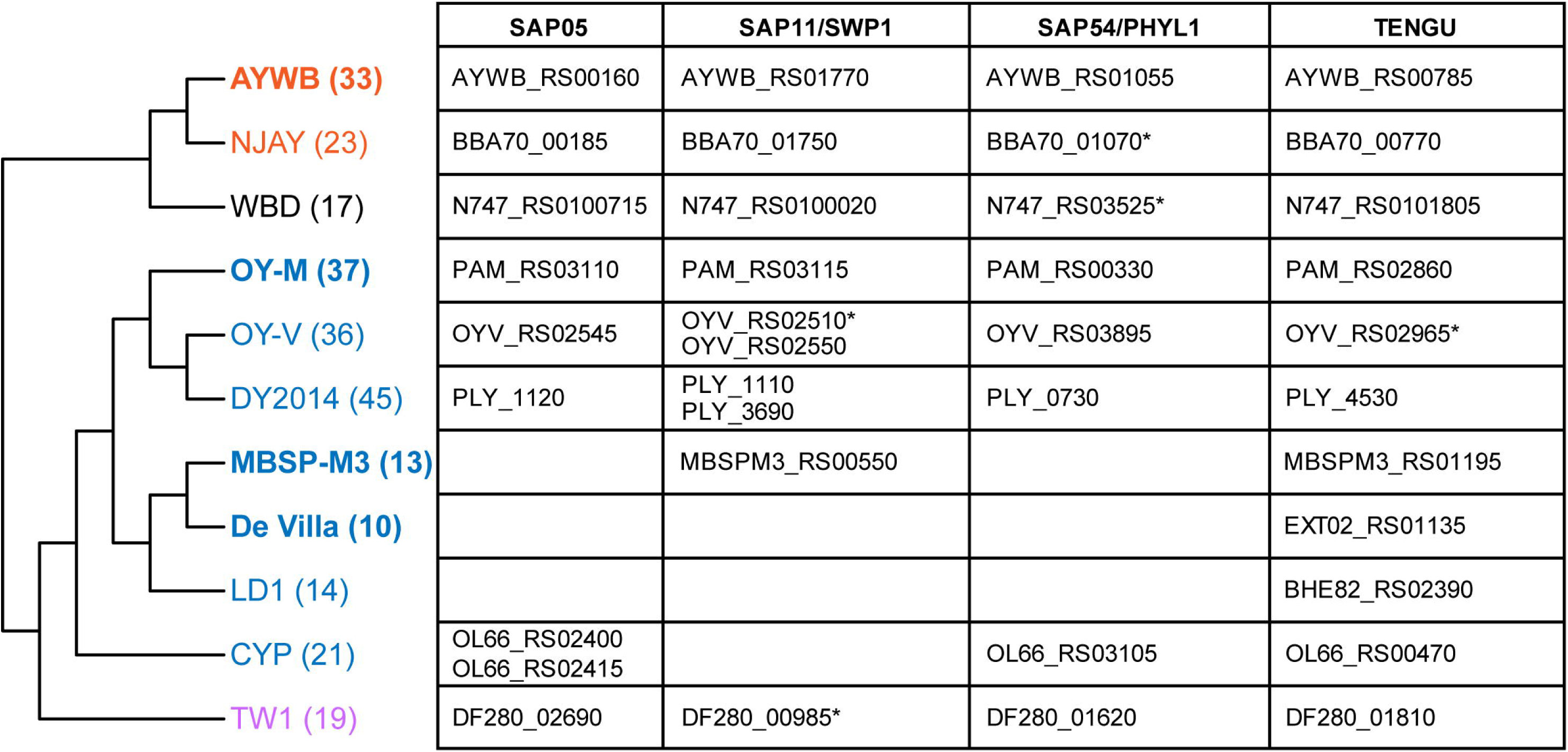
Phylogenetic distribution of putative effector genes. The cladogram that illustrates the phylogenetic relationships among these strains is based on Figure 1B (i.e., maximum likelihood phylogeny based on 303 shared single-copy genes). The number in parentheses after the strain name indicates the number of genes that encode putative secreted proteins in each genome (including putative pseudogenes). The strains with the complete genome sequences available are highlighted in bold. On the right-hand side, the table lists the locus tags of homologs for those four phytoplasma effectors that have been characterized. Those locus tags with an asterisk (*) were annotated as putative pseudogenes.

For those 16SrI phytoplasma effectors that have been characterized experimentally, namely SAP05 (Gamboa et al., 2019; Huang and Hogenhout, 2019), SAP11/SWP1 (Bai et al., 2009; Sugio et al., 2011a; Lu et al., 2014; Chang et al., 2018; Wang et al. 2018b; Wang et al. 2018c), SAP54/PHYL1 (MacLean et al., 2011; Maejima et al., 2014; Orlovskis and Hogenhout, 2016), and TENGU (Hoshi et al., 2009; Sugawara et al., 2013; Minato et al., 2014), only TENGU is conserved among all strains. Although some cases of gene absence may be artifacts of incomplete genome assemblies, the strain De Villa (which has the complete chromosomal sequence available) lacks all other three effector genes, indicating that none of those three effectors is essential.

The variable pattern of effector gene presence and absence may be linked to the molecular evolution of potential mobile units (PMU) in these phytoplasma genomes (Bai et al., 2006). These mobile genetic elements were known to facilitate horizontal gene transfers between either closely or distantly related strains and may be gained or lost rapidly (Chung et al., 2013; Ku et al., 2013; Orlovskis et al., 2017; Cho et al., 2019b; Seruga Music et al., 2019). When we examined the chromosomal locations of those four effector genes, TENGU is not associated with PMUs, while the other three are. Finally, as noted in our previous study (Cho et al., 2019b), DY2014 and OY-V both have two copies of SAP11 homologs. One copy is located within a 16SrI-B type PMU, suggesting vertical inheritance, while the other copy is located within a 16SrI-A type PMU, suggesting horizontal acquisition. These observations provided further support to the importance of PMUs in phytoplasma effector evolution.

## Conclusion

By quantifying the genetic divergence among a group of closely-related phytoplasmas, this work demonstrated a discontinuity in their similarities of genomic content and homologous sequences. The results suggested that these phytoplasmas could be classified into multiple distinct taxonomic units equivalent to species in other bacteria (Jain et al., 2018). Importantly, the widely used classification system for phytoplasmas that is based on the sequencing and RFLP analysis of their 16S rRNA genes does not provide sufficient resolution or accuracy. Rather, it is important to incorporate genomic information in the future revisions of phytoplasma taxonomy. A previous proposal of using 95% ANI as a cutoff to delineate bacterial species (Jain et al., 2018) appears to work well and warrants further consideration to be adopted. In recognition of the difficulties involved in genomic studies of uncultivated bacteria, this work also demonstrated the feasibility of using genome analysis to develop cost-effective MLSA markers, which will be useful for practical purposes. For future directions, it is critical to improve the taxon sampling of phytoplasma genomes strategically, such that similar approaches may be expand to genus-wide analysis to facilitate the studies of these important plant pathogens. Moreover, continuing effort in establishing axenic culture is critical for the study of these important plant pathogens. In the absence of axenic culture, it is important that the reference strains, particularly those with provisional species status, are made available to the scientific community, preferably in the form of a centralized micropropagation collection as suggested by the international phytoplasma working team (The IRPCM Phytoplasma/Spiroplasma Working Team, 2004). If live culture within plant or insect hosts is not possible, then the DNA samples should be made available. For genome sequencing projects, the raw sequencing results should be made publicly available together with the assembled genome sequences at the time of publication, such that the assemblies could be validated independently. The TW1 genome (Town et al., 2018) provided a good example to illustrate the importance of this point. Although the publication of an erroneous genome sequence is unfortunate, the raw reads made available by those authors allowed us to investigate and identify the issues; such openness is appreciated and important for science to move forward.

## Acknowledgments

We thank Ms. Mei-Jane Fang in the Genomic Technology Core Facility (Institute of Plant and Microbial Biology, Academia Sinica) for providing Sanger sequencing service. This manuscript was posted as a preprint at bioRxiv (https://doi.org/10.1101/2020.01.31.928135). Comments received during the peer review process greatly improved this work. This manuscript is not meant to be a comprehensive review of all the available literature on phytoplasma classification; as such we apologize if we did not cite all publications on this topic in the manuscript.

## Author Contribution

S-TC, H-JK, and C-HK performed the experiments and analyzed the data. WH and SAH provided intellectual input. S-TC prepared the figures and supplementary materials. S-TC and C-HK wrote the manuscript. SAH edited the manuscript. C-HK designed the experiments, acquired the funding, and supervised the project.

## Supplementary Materials

Supplementary Figure S1. Maximum likelihood phylogenies. Based on Figure 1, the TW1 sequences were excluded prior to multiple sequence alignment. The numbers on the internal branches indicate the level of bootstrap support based on 1,000 resampling; only values >=60% are shown. (A) 16S rRNA genes. The strain name, locus tag of the 16S rRNA gene (in parentheses), and the 16SrI subgroup classification (in square brackets) are labeled. A question mark ‘?’ in the subgroup classification indicates that the sequence is classified as a new subgroup, the existing subgroup with the highest similarity is provided. (B) The 303 single-copy coding genes shared by all strains. (C) 16S rRNA gene and the five markers developed in this study; see Table 2 for detailed information.

Supplementary Figure S2. Multiple sequence alignments of selected marker genes. The primer sites are highlighted in orange. Shades of blue colors in the alignment indicate the levels of sequence conservation.

Supplementary Figure S3. Distribution of substitution rates among those single-copy genes shared by the 11 phytoplasma genomes analyzed. The synonymous substitution rate (Ks) and non-synonymous substitution rate (Ka) were calculated based on the homologs found in AYWB and MBSP-M3. Among those 303 single-copy genes, nine were annotated as pseudogenes and were omitted from the substitution rate calculation.

Supplementary Table S1. Pairwise nucleotide sequence identity of the molecular markers used in the phylogenetic inference. Numbers in the diagonal cells indicate the length of each unaligned sequence.

Supplementary Table S2. List of the 303 single-copy genes shared by all of the 11 phytoplasma genomes analyzed in this study. The gene identity, genomic location, and functional annotation of these genes are provided based on the information from AYWB. The substitution rate estimates are based on the comparison between AYWB and MBSP-M3.

Supplementary Table S3. List of the putative effector genes. Each gene is uniquely identified by its locus_tag. The “Cluster_id” field provides the assignment of homologous gene cluster; genes sharing the same Cluster_id are considered as homologs. Note that SAP11/SWP1 homologs are separated into two clusters (i.e., 503 and 598) due to high levels of sequence divergence. Additionally, one SAP05 homolog is in a separate cluster (i.e., 1117) due to truncation. The gene names and product description are based on existing GenBank annotation; additional information (e.g., homology information with known effector or pseudogene status) are provided in the “Note” field.

