## Supplementary figures and images for "Species boundaries and molecular markers for the classification of 16SrI phytoplasmas inferred by genome analysis"

### Supplementary Figure S1

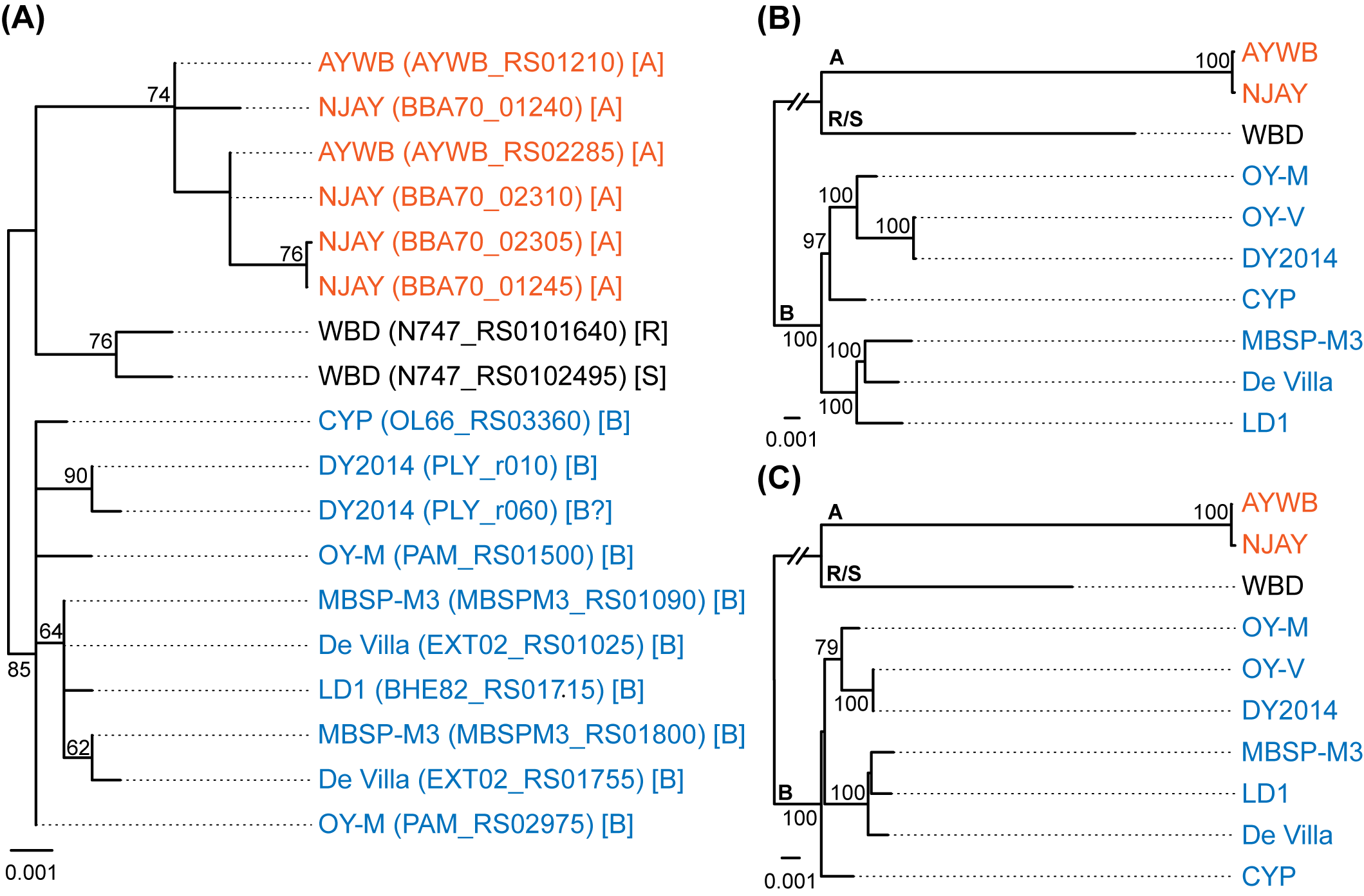

### Supplementary Figure S3

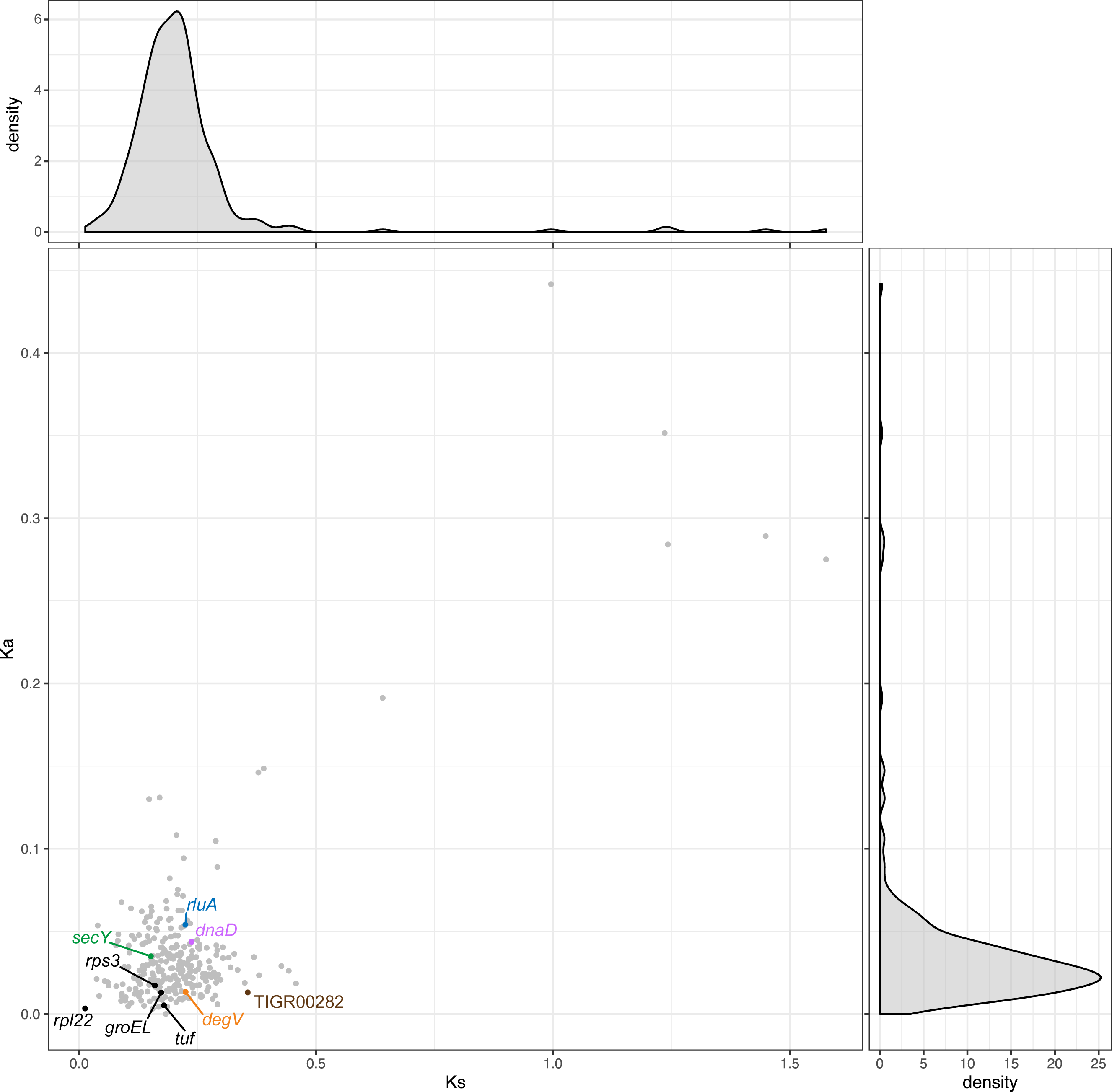
