## Supplementary Figure S2 for "Species boundaries and molecular markers for the classification of 16SrI phytoplasmas inferred by genome analysis"

### (A) replication initiation protein DnaD

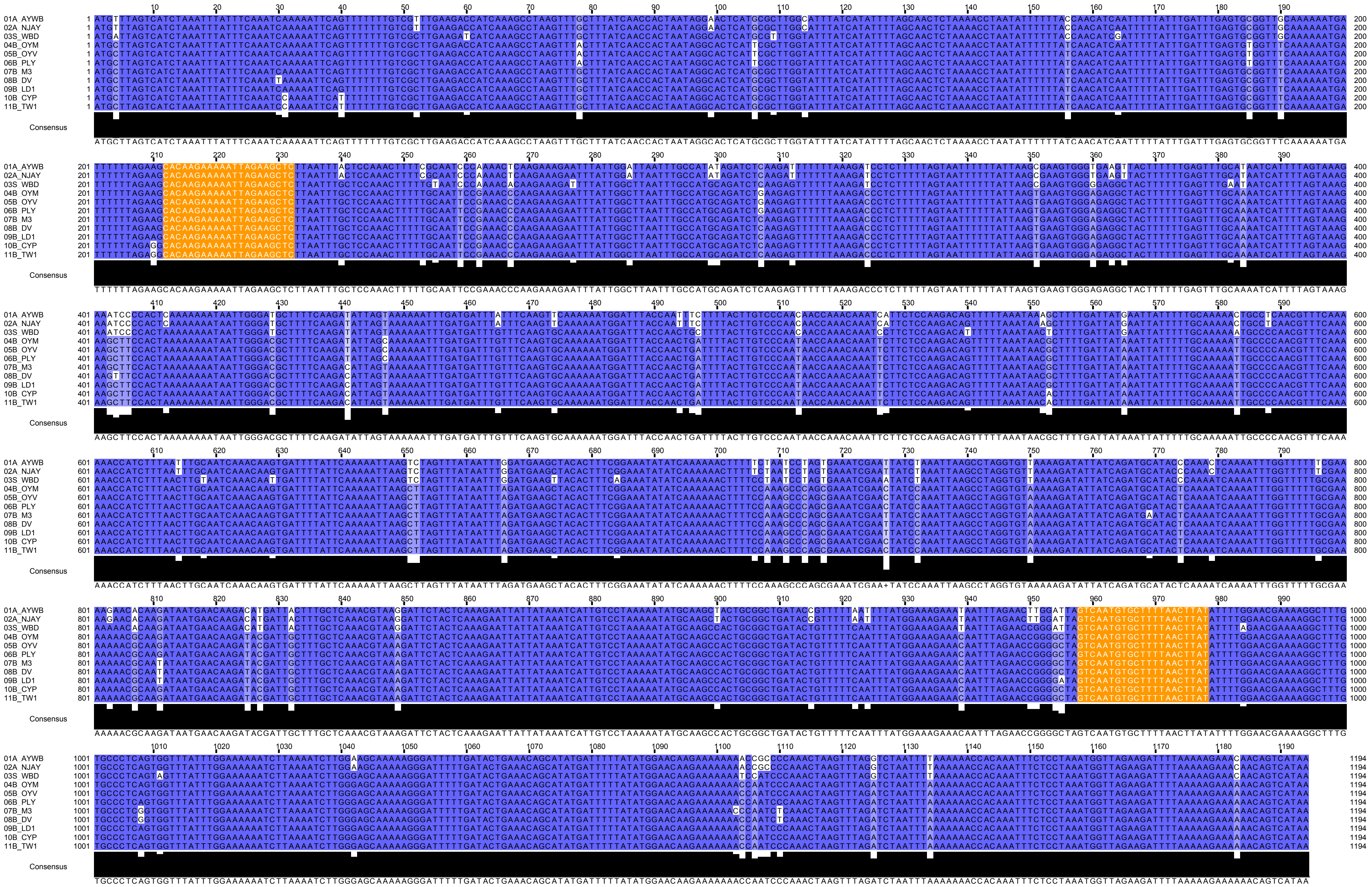

### (B) DegV family protein

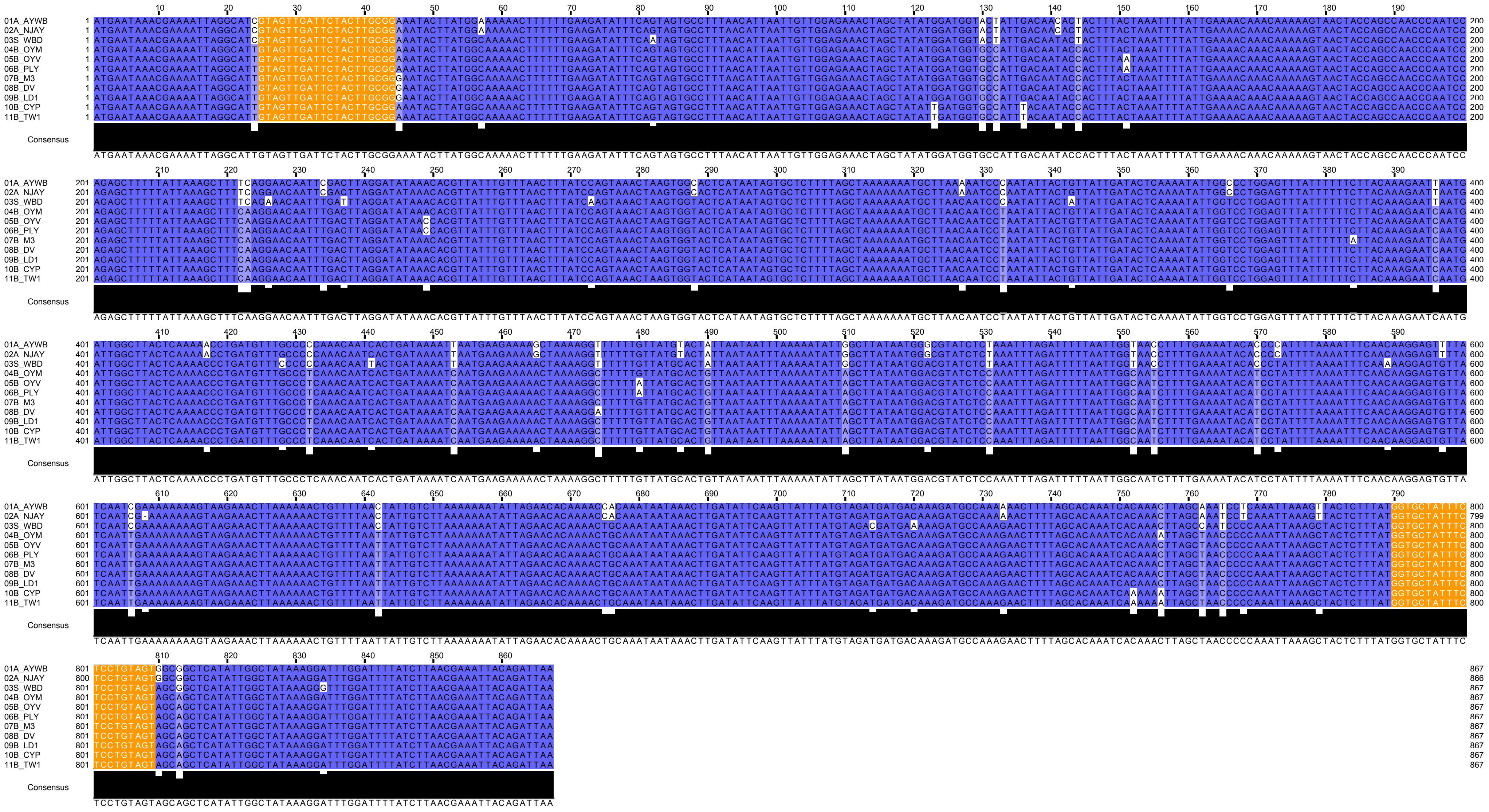

### (C) TIGR00282 family metallophosphoesterase

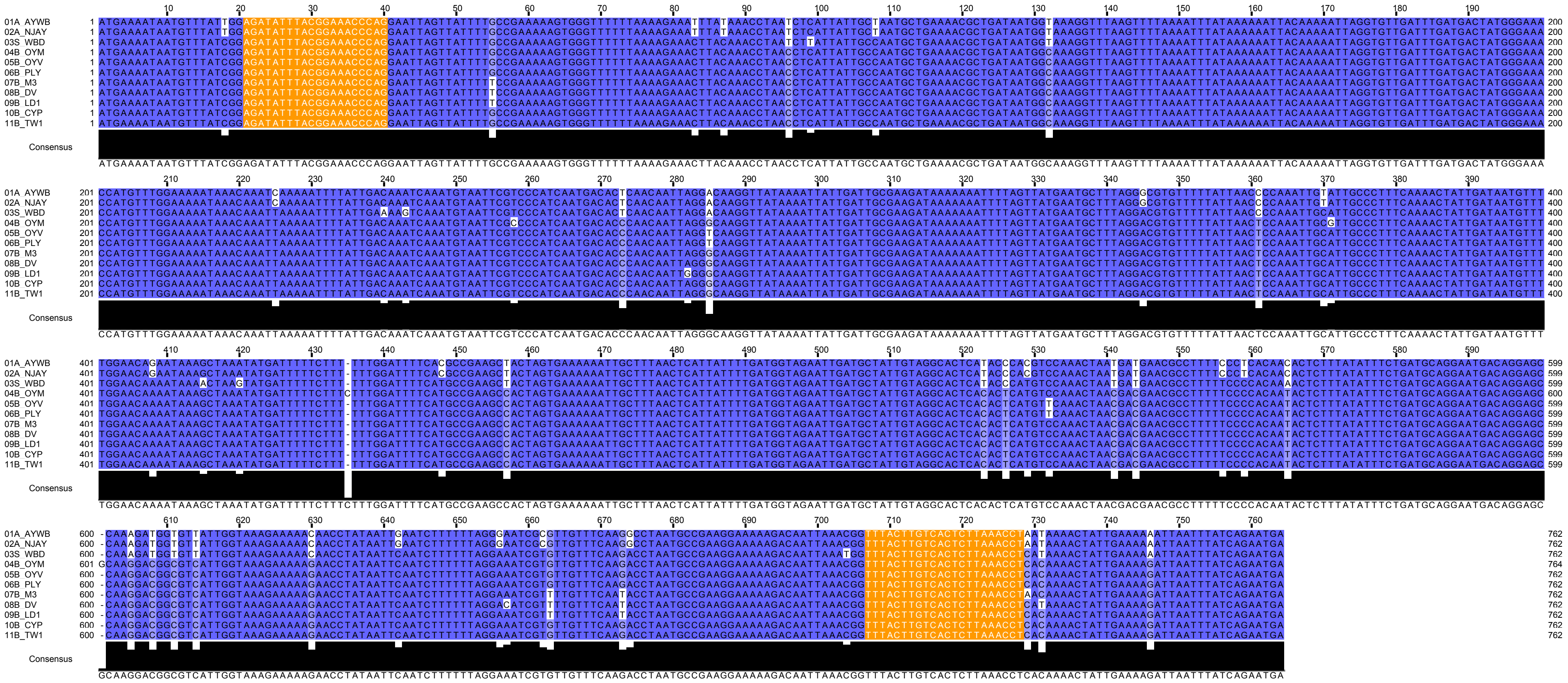

### (D) preprotein translocase subunit SecY

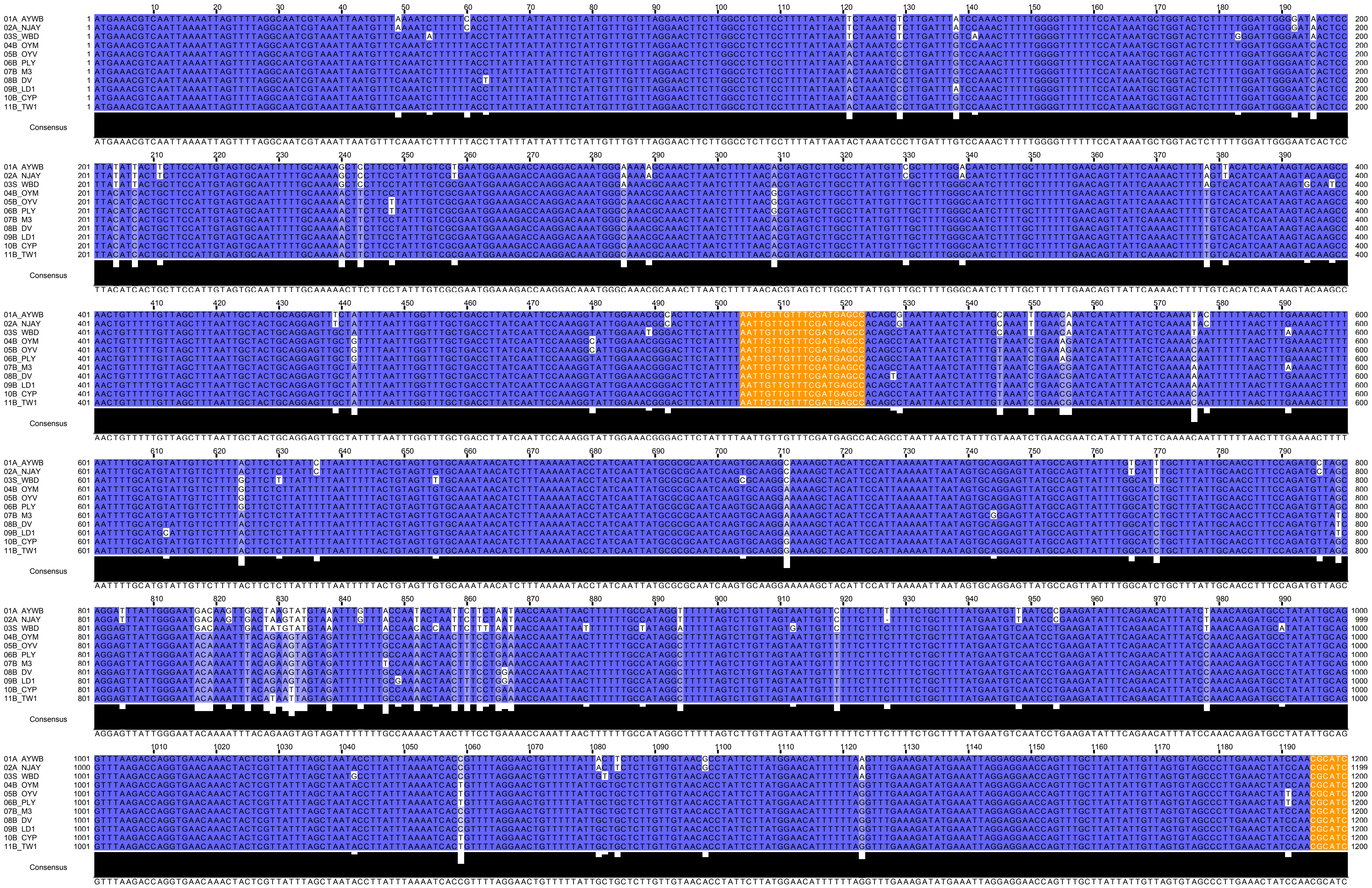

| | | 1210 1220 1230 1240 | |
|---|---|---|---|
| 01A_AYWB | 1201 | AAAGCTACTGCCAACAAAAAAGAATATCAAAAATTATTTTAA | 1242 |
| 02A_NJAY | 1200 | AAAGCTACTGCCAACAAAAAAGAATATCAAAAATTATTTTAA | 1241 |
| 03S_WBD | 1201 | AAAGCTACTGCCAACAAAAAAGAATATCAAAAATTATTTTAA | 1242 |
| 04B_OYM | 1201 | AAAGCTACTGCCAACAAAAAAGAATATCAAAAATTATTTTAA | 1242 |
| 05B_OYV | 1201 | AAAGCTACTGCCAACAAAAAAGAATATCAAAAATTATTTTAA | 1242 |
| 06B_PLY | 1201 | AAAGCTACTGCCAACAAAAAAGAATATCAAAAATTATTTTAA | 1242 |
| 07B_M3 | 1201 | AAAGCTACTGCCAACAAAAAAGAATATCAAAAATTATTTTAA | 1242 |
| 08B_DV | 1201 | AAAGCTACTGCCAACAAAAAAGAATATCAAAAATTATTTTAA | 1242 |
| 09B_LD1 | 1201 | AAAGCTACTGCCAACAAAAAAGAATATCAAAAATTATTTTAA | 1242 |
| 10B_CYP | 1201 | AAAGCTACTGCCAACAAAAAAGAATATCAAAAATTATTTTAA | 1242 |
| 11B_TW1 | 1201 | AAAGCTACTGCCAACAAAAAAGAATATCAAAAATTATTTTAA | 1242 |

Consensus

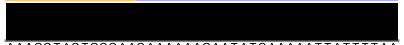

AAAGCTACTGCCAACAAAAAAGAATATCAAAAATTATTTTAA

### (E) RluA family pseudouridine synthase

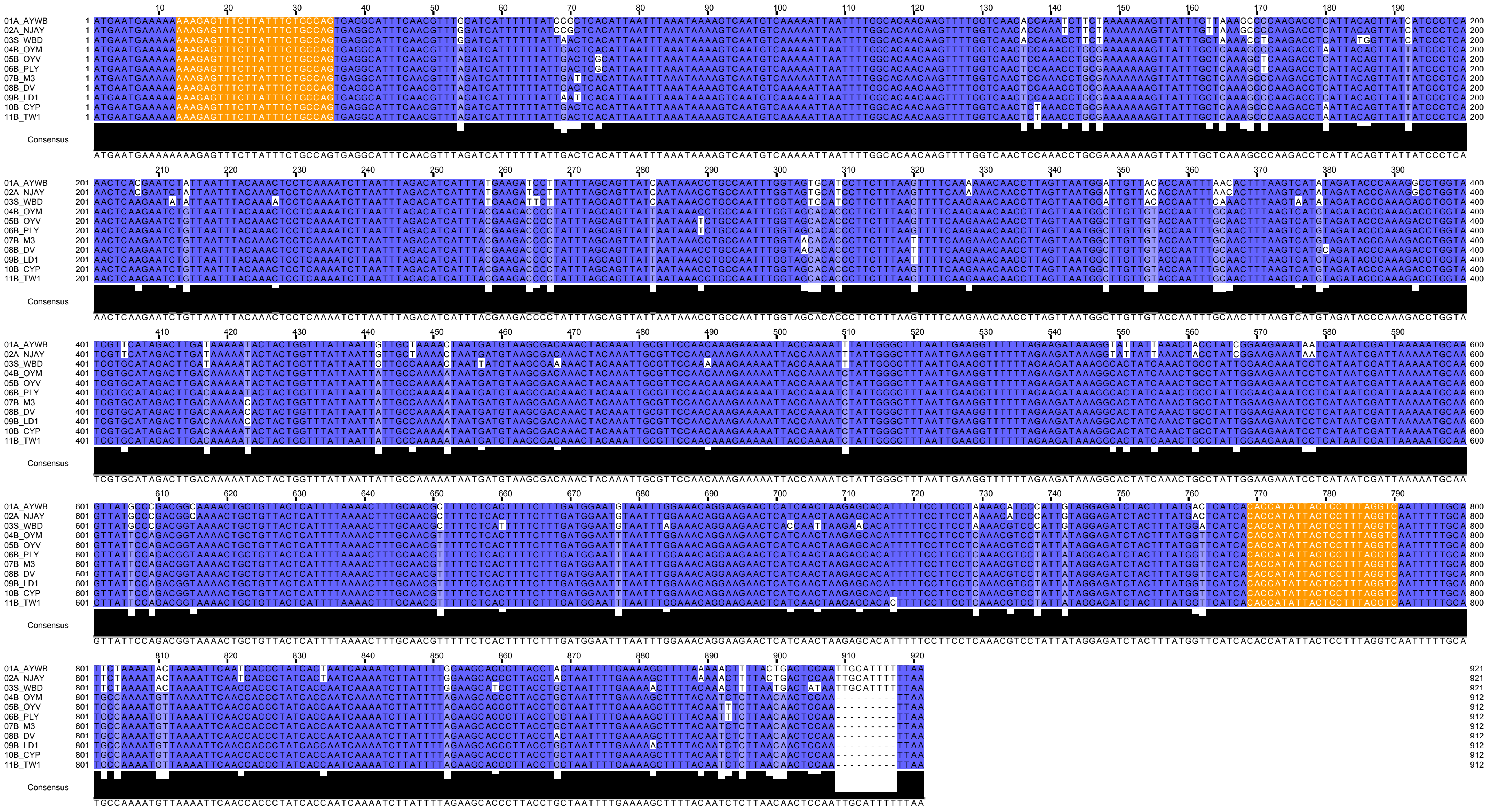

### (R) 16S ribosomal RNA

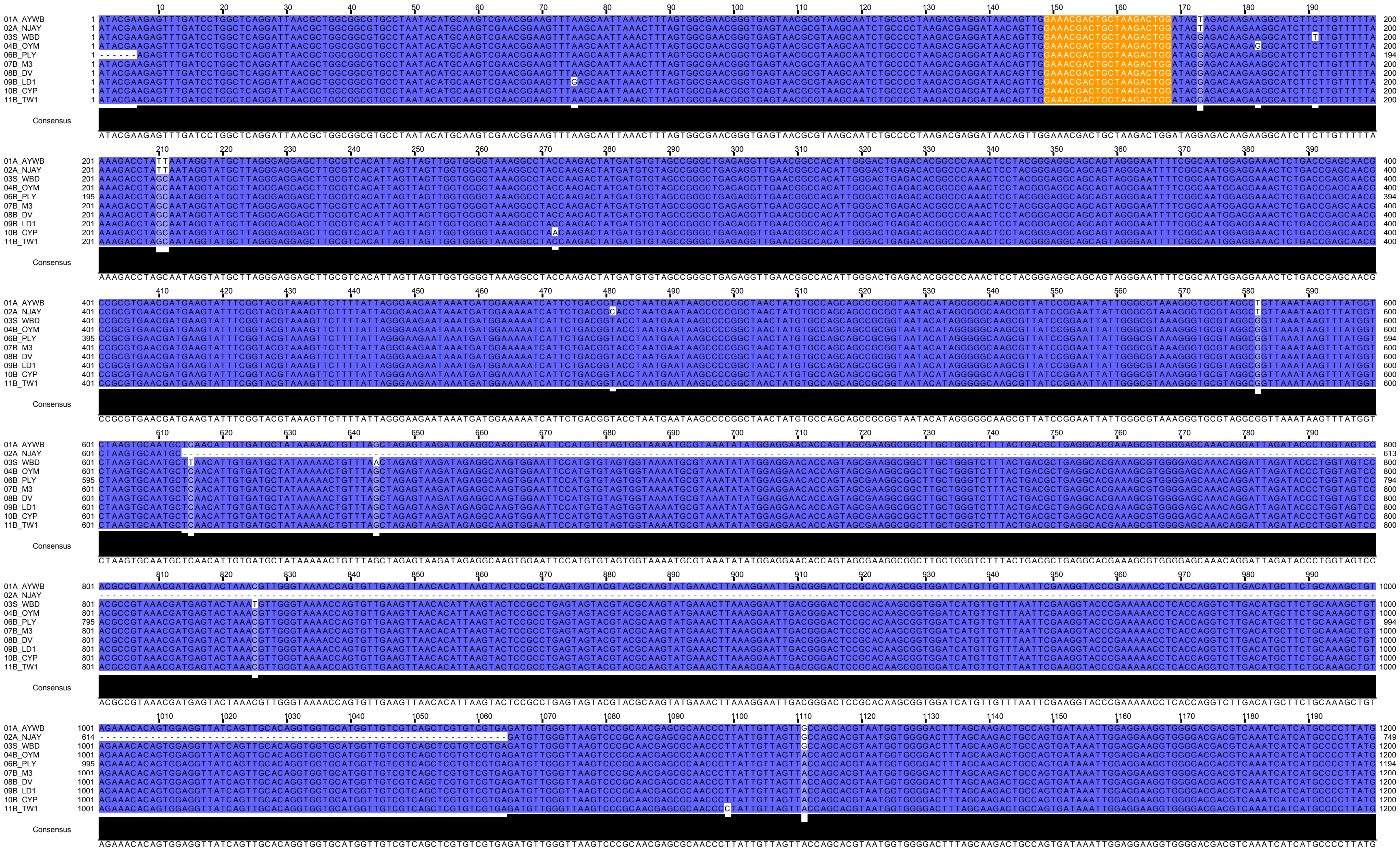

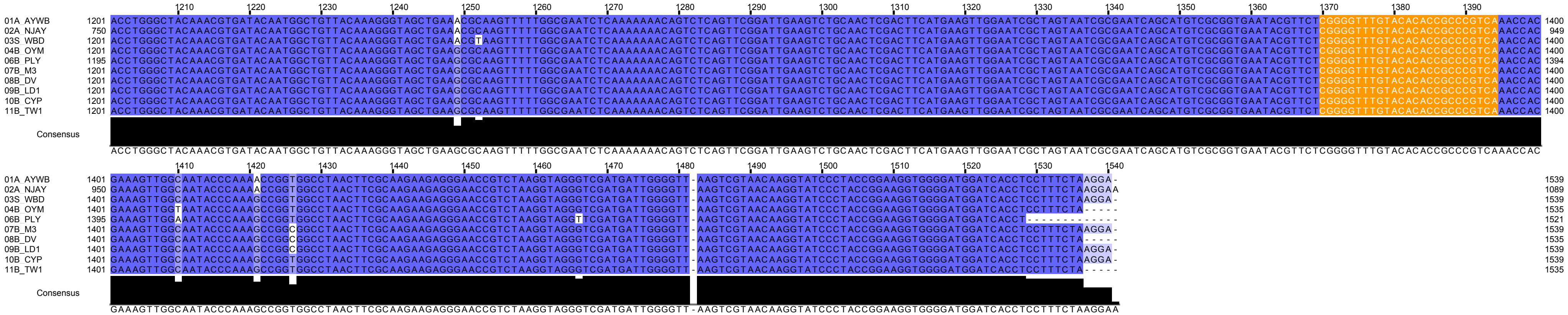
1210 1220 1230 1240 1250 1260 1270 1280 1290 1300 1310 1320 1330 1340 1350 1360 1370 1380 1390
01A AYWB 1201 ACCTGGGCTACAAACGTGATACAATGGCTGTTACAAAGGGTAGCTGAAACGCAAGTTTTTGGCGAATCTCAAAAAAACAGTCTCAGTTCGGATTGAAGTCTGCAACTCGACTTCATGAAGTTGGAATCGCTAGTAATCGCGAATCAGCATGTCGCGGTGAATACGTTCTCGGGGTTTGTACACACCGCCCGTCAAACCAC 1400
02A NJAY 750 ACCTGGGCTACAAACGTGATACAATGGCTGTTACAAAGGGTAGCTGAAACGCAAGTTTTTGGCGAATCTCAAAAAAACAGTCTCAGTTCGGATTGAAGTCTGCAACTCGACTTCATGAAGTTGGAATCGCTAGTAATCGCGAATCAGCATGTCGCGGTGAATACGTTCTCGGGGTTTGTACACACCGCCCGTCAAACCAC 949
03S WBD 1201 ACCTGGGCTACAAACGTGATACAATGGCTGTTACAAAGGGTAGCTGAAACGTAAGTTTTTGGCGAATCTCAAAAAAACAGTCTCAGTTCGGATTGAAGTCTGCAACTCGACTTCATGAAGTTGGAATCGCTAGTAATCGCGAATCAGCATGTCGCGGTGAATACGTTCTCGGGGTTTGTACACACCGCCCGTCAAACCAC 1400
04B OYM 1201 ACCTGGGCTACAAACGTGATACAATGGCTGTTACAAAGGGTAGCTGAAGCGCAAGTTTTTGGCGAATCTCAAAAAAACAGTCTCAGTTCGGATTGAAGTCTGCAACTCGACTTCATGAAGTTGGAATCGCTAGTAATCGCGAATCAGCATGTCGCGGTGAATACGTTCTCGGGGTTTGTACACACCGCCCGTCAAACCAC 1400
06B PLY 1195 ACCTGGGCTACAAACGTGATACAATGGCTGTTACAAAGGGTAGCTGAAGCGCAAGTTTTTGGCGAATCTCAAAAAAACAGTCTCAGTTCGGATTGAAGTCTGCAACTCGACTTCATGAAGTTGGAATCGCTAGTAATCGCGAATCAGCATGTCGCGGTGAATACGTTCTCGGGGTTTGTACACACCGCCCGTCAAACCAC 1394
07B_M3 1201 ACCTGGGCTACAAACGTGATACAATGGCTGTTACAAAGGGTAGCTGAAGCGCAAGTTTTTGGCGAATCTCAAAAAAACAGTCTCAGTTCGGATTGAAGTCTGCAACTCGACTTCATGAAGTTGGAATCGCTAGTAATCGCGAATCAGCATGTCGCGGTGAATACGTTCTCGGGGTTTGTACACACCGCCCGTCAAACCAC 1400
08B_DV 1201 ACCTGGGCTACAAACGTGATACAATGGCTGTTACAAAGGGTAGCTGAAGCGCAAGTTTTTGGCGAATCTCAAAAAAACAGTCTCAGTTCGGATTGAAGTCTGCAACTCGACTTCATGAAGTTGGAATCGCTAGTAATCGCGAATCAGCATGTCGCGGTGAATACGTTCTCGGGGTTTGTACACACCGCCCGTCAAACCAC 1400
09B_LD1 1201 ACCTGGGCTACAAACGTGATACAATGGCTGTTACAAAGGGTAGCTGAAGCGCAAGTTTTTGGCGAATCTCAAAAAAACAGTCTCAGTTCGGATTGAAGTCTGCAACTCGACTTCATGAAGTTGGAATCGCTAGTAATCGCGAATCAGCATGTCGCGGTGAATACGTTCTCGGGGTTTGTACACACCGCCCGTCAAACCAC 1400
10B CYP 1201 ACCTGGGCTACAAACGTGATACAATGGCTGTTACAAAGGGTAGCTGAAGCGCAAGTTTTTGGCGAATCTCAAAAAAACAGTCTCAGTTCGGATTGAAGTCTGCAACTCGACTTCATGAAGTTGGAATCGCTAGTAATCGCGAATCAGCATGTCGCGGTGAATACGTTCTCGGGGTTTGTACACACCGCCCGTCAAACCAC 1400
11B_TW1 1201 ACCTGGGCTACAAACGTGATACAATGGCTGTTACAAAGGGTAGCTGAAGCGCAAGTTTTTGGCGAATCTCAAAAAAACAGTCTCAGTTCGGATTGAAGTCTGCAACTCGACTTCATGAAGTTGGAATCGCTAGTAATCGCGAATCAGCATGTCGCGGTGAATACGTTCTCGGGGTTTGTACACACCGCCCGTCAAACCAC 1400
Consensus
ACCTGGGCTACAAACGTGATACAATGGCTGTTACAAAGGGTAGCTGAAGCGCAAGTTTTTGGCGAATCTCAAAAAAACAGTCTCAGTTCGGATTGAAGTCTGCAACTCGACTTCATGAAGTTGGAATCGCTAGTAATCGCGAATCAGCATGTCGCGGTGAATACGTTCTCGGGGTTTGTACACACCGCCCGTCAAACCAC
1410 1420 1430 1440 1450 1460 1470 1480 1490 1500 1510 1520 1530 1540
01A AYWB 1401 GAAAGTTGGCAATACCCAAAACCGGTGGCCTAACTTCGCAAGAAGAGGGAACCGTCTAAGGTAGGGTCGATGATTGGGGTT-AAGTCGTAACAAGGTATCCCTACCGGAAGGTGGGGATGGATCACCTCCTTTCTAAGGA- 1539
02A NJAY 950 GAAAGTTGGCAATACCCAAAACCGGTGGCCTAACTTCGCAAGAAGAGGGAACCGTCTAAGGTAGGGTCGATGATTGGGGTT-AAGTCGTAACAAGGTATCCCTACCGGAAGGTGGGGATGGATCACCTCCTTTCTAAGGAA 1089
03S WBD 1401 GAAAGTTGGCAATACCCAAAGCCGGTGGCCTAACTTCGCAAGAAGAGGGAACCGTCTAAGGTAGGGTCGATGATTGGGGTT-AAGTCGTAACAAGGTATCCCTACCGGAAGGTGGGGATGGATCACCTCCTTTCTAAGGA- 1539
04B OYM 1401 GAAAGTTGGTAATACCCAAAGCCGGTGGCCTAACTTCGCAAGAAGAGGGAACCGTCTAAGGTAGGGTCGATGATTGGGGTT-AAGTCGTAACAAGGTATCCCTACCGGAAGGTGGGGATGGATCACCTCCTTTCTA----- 1535
06B PLY 1395 GAAAGTTGGAAATACCCAAAGCCGGTGGCCTAACTTCGCAAGAAGAGGGAACCGTCTAAGGTAGGTTCGATGATTGGGGTT-AAGTCGTAACAAGGTATCCCTACCGGAAGGTGGGGATGGATCACCT------------- 1521
07B_M3 1401 GAAAGTTGGCAATACCCAAAGCCGGCGGCCTAACTTCGCAAGAAGAGGGAACCGTCTAAGGTAGGGTCGATGATTGGGGTT-AAGTCGTAACAAGGTATCCCTACCGGAAGGTGGGGATGGATCACCTCCTTTCTAAGGA- 1539
08B_DV 1401 GAAAGTTGGCAATACCCAAAGCCGGCGGCCTAACTTCGCAAGAAGAGGGAACCGTCTAAGGTAGGGTCGATGATTGGGGTT-AAGTCGTAACAAGGTATCCCTACCGGAAGGTGGGGATGGATCACCTCCTTTCTA----- 1535
09B_LD1 1401 GAAAGTTGGCAATACCCAAAGCCGGCGGCCTAACTTCGCAAGAAGAGGGAACCGTCTAAGGTAGGGTCGATGATTGGGGTT-AAGTCGTAACAAGGTATCCCTACCGGAAGGTGGGGATGGATCACCTCCTTTCTAAGGA- 1539
10B CYP 1401 GAAAGTTGGCAATACCCAAAGCCGGTGGCCTAACTTCGCAAGAAGAGGGAACCGTCTAAGGTAGGGTCGATGATTGGGGTT-AAGTCGTAACAAGGTATCCCTACCGGAAGGTGGGGATGGATCACCTCCTTTCTAAGGA- 1539
11B_TW1 1401 GAAAGTTGGCAATACCCAAAGCCGGTGGCCTAACTTCGCAAGAAGAGGGAACCGTCTAAGGTAGGGTCGATGATTGGGGTT-AAGTCGTAACAAGGTATCCCTACCGGAAGGTGGGGATGGATCACCTCCTTTCTA----- 1535
Consensus
GAAAGTTGGCAATACCCAAAGCCGGTGGCCTAACTTCGCAAGAAGAGGGAACCGTCTAAGGTAGGGTCGATGATTGGGGTT-AAGTCGTAACAAGGTATCCCTACCGGAAGGTGGGGATGGATCACCTCCTTTCTAAGGAA
